# Non-Uniformity of Projection Distributions Attenuates Resolution in Cryo-EM

**DOI:** 10.1101/635938

**Authors:** Philip R. Baldwin, Dmitry Lyumkis

## Abstract

Virtually every single-particle cryo-EM experiment currently suffers from specimen adherence to the air-water interface, leading to a non-uniform distribution in the set of projection views. Whereas it is well accepted that uniform projection distributions can lead to high-resolution reconstructions, non-uniform (anisotropic) distributions can negatively affect map quality, elongate structural features, and in some cases, prohibit interpretation altogether. Although some consequences of non-uniform sampling have been described qualitatively, we know little about how sampling quantitatively affects resolution in cryo-EM, especially given the numerous different projection schemes that can arise in experimental situations. Here, we show how inhomogeneity in any projection distribution scheme attenuates the global Fourier Shell Correlation (FSC) in relation to the number of particles and a single geometrical parameter, which we term the sampling compensation factor (SCF). The reciprocal of the SCF is defined as the average over Fourier shells of the reciprocal of the per-particle sampling and normalized to unity for uniform distributions. The SCF therefore ranges from one to zero, with values close to the latter implying large regions of poorly sampled or completely missing data in Fourier space. Using two synthetic test cases, influenza hemagglutinin and human apoferritin, we demonstrate how any amount of sampling inhomogeneity always attenuates the FSC compared to a uniform distribution. We advocate quantitative evaluation of the SCF criterion to approximate the effect of non-uniform sampling on resolution within experimental single-particle cryo-EM reconstructions.

## Introduction

Single-particle cryo-electron microscopy (cryo-EM) has gained increasing popularity for structural analysis of macromolecules and macromolecular assemblies. Numerous technical advances have contributed to improvements in resolution [1-3], throughput [4], and overall usability of the approaches, leading to a wealth of novel insights pertaining to macromolecular structure and function [5]. Although many steps in the single-particle workflow are becoming more streamlined and automated, a principal remaining challenge pertains to problems resulting from non-uniform projection distributions contributing to reconstructed density maps.

Non-uniformity in the distribution of projection orientations recorded in a single-particle imaging experiment originates from adherence of the specimen to one of two interfaces (top or bottom) of the grid. The interfaces, which could be air-water or support-water (e.g. thin carbon), cause specimens to stick in one of several “preferential orientations”. It is now clear that virtually every specimen prepared for single-particle imaging using conventional blotting techniques adopts a preferential orientation on cryo-EM grids [6]. The reason for this is that macromolecules, which continuously undergo rapid thermal motion, adhere to interfaces on a time scale that is orders of magnitude shorter than the time to blot off excess sample. Recent inkjet dispensing technologies have ameliorated some of the effects of preferential specimen orientation by attempting to out-run sample adherence to interfaces and by minimizing the amount of time between sample application and plunging into liquid ethane [7]. However, such devices do not yet eliminate preferential orientation in its entirety and depend heavily on high sample concentration. Furthermore, the increase in interest in specimen supports, like graphene [8, 9], which also cause preferential orientation, indicates that the effects of non-uniform sampling on final reconstructions will remain problematic for many single-particle experiments.

Numerous approaches have been devised to estimate the quality of angular distributions and their effects on a reconstructed density. These ideas are primarily developed in conjunction with some anisotropic measure. One measure derives from the application of a 3D point spread function to estimate the strength of signal above some significance criterion, in all directions of the 3D Fourier transform [10]. In another approach, the 3D spectral signal-to noise ratio (SSNR) is used to define directional resolution differences [11], with the SSNR bearing a direct relationship to the Fourier Shell Correlation (FSC), the conventional means for measuring resolution in single-particle cryo-EM[12]. Multiple groups also described the use of conical FSCs to evaluate anisotropic resolution for tomographic reconstructions [13, 14], as well as our and others’ work on evaluating anisotropic resolution in single-particle analysis [15, 16]. More recently, the “efficiency” metric [17] was introduced to characterize an orientation distribution, based on the observed relationship between orientation distribution and experimental resolution. We proposed that an evaluation of anisotropy in cryo-EM experiments should be standard for every cryo-EM reconstruction [18].

The consequences of sampling non-uniformity on a reconstructed density map can vary and depend on the extent and distribution of projection views. In many experimental cases, one might see a few distinct preferential orientations across the Euler distribution profile, but the resulting map may look reasonable, and is readily interpretable with an atomic model. In the more severe cases, an anisotropic distribution may lead to apparent elongation of structural features within the map. In such cases, the interpretation of the map may be affected, sometimes severely, due to the appearance of artefactual density parallel to the dominant view [16]. In the most severe cases, structure determination may be stifled altogether. Some hallmarks of pathologically anisotropic distributions include inflated Fourier Shell Correlation (FSC) curves, elongated features beyond interpretability, an inability to converge on a final structure, and/or the appearance of false positive orientations in the course of refinement [16]. All these factors can reinforce problems in the density. One interesting observation was that anisotropic orientation distributions lead to an increase in the temperature factor associated with the data, thereby also affecting global resolution [17]. However, a derivation from standard models has not been established.

While different measures have been introduced to evaluate the effect of anisotropic distributions on directional viewings of the reconstructed density map, the effect of sampling on global resolution has largely been neglected. Furthermore, there remains no systematic, quantitative study of the effects of inhomogeneous projection distributions on cryo-EM reconstructions. Here, we examine the relationship between non-uniform angular sampling and global resolution, as measured using conventional analyses in cryo-EM. A major conclusion from our work is that *any* inhomogeneity, and especially missing information in Fourier space, directly attenuates global resolution in 3D reconstructions, and thus impedes the single-particle experiment.

## Section 1. Summary of the major findings

Given a set of projection views, we develop an assessment of the quality of the sampling. We chose this assessment based on the expected effect on the spectral signal to noise ratio (SSNR) defined through the FSC. We show that the angular average of the reciprocal of the sampling forms a quantity whose reciprocal attenuates the SSNR, if we consider the other aspects of the problem associated with the overall experimental envelope to be held constant. More specifically, we argue:

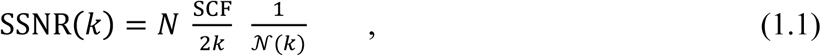

where *N* is the number of particles, *k* is spatial frequency, 𝒩(*k*) is a noise-to-signal power, and SCF is what we term the “sampling compensation factor” and is defined to be

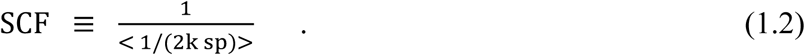

Here, <·> means the average in Fourier space over the nonzero values of shells at (approximately) fixed spatial frequency, *k*, and sp is the amount of sampling per-particle, determined from the Euler angle assignments. Notably, one must compensate for the geometry of the sampling to correctly estimate the SSNR: hence the name, “sampling compensation factor”. One notes from (1.1) that the number of particles necessary to perform a reconstruction also depends inversely on the SCF, with smaller SCFs requiring larger numbers of particles.

In section 2, we derive all the formulae relating sampling to SSNR, including the case with missing data, which requires special handling. In section 3, we derive analytical solutions to the sampling and SCF for a variety of different cases. In section 4, we discuss the linear dependence of the SSNR on N, as well as estimating the number of particles to perform a reconstruction. In section 5, we show the correspondence between the proposed decrement of signal based on sampling and the actual decrement in the SSNR when reconstructions are performed for two different proteins.

## Section 2. Decrement of SSNR due to sampling inhomogeneity

In this section, we derive Eq (1.1), which provides an expression for the SSNR where all the aspects of the sampling have been incorporated specifically into two parameters: the number of particles, and a single geometrical factor. We assume that the effects of the microscope and the effects of the noise can be approximately decoupled, in a manner that has otherwise been typically assumed in the literature [19-21]. In section 2.1, we first consider the cases where the voxels in 3D Fourier space are completely measured and derive the SSNR relationship, Eqs (1.1) and (1.2), which is the main result of this paper. In section 2.2, we extend these derivations to cases when there is missing data, by which we arrive at the adjusted formulae for resolution (2.30). We refer to other sources, as necessary (Sorzano, [21] and Penczek [19]), for more detail on the aspects that are not central to the derivations given here.

### 2.1 Derivation of the Sampling Compensation Factor (SCF)

The generally accepted understanding of 2D projection data after orientation assignment in cryo-EM single-particle analysis is given by:

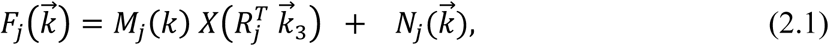

Here 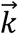 is a point in 2D Fourier space as measured on the projection *j*, where the projection *j* has data 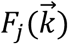 on the 2D grid point labeled by 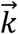 (see Figure 1). This is the usual Fourier space description of a “single particle”. Eq (2.1) is our statement of the projection slice theorem: the measured data should be a slice out of the true 3D map, *X*, but that has been modified in the microscope by a transfer function, *M*_*j*_ (k) and corrupted by 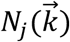, which is identically distributed noise with mean zero and a variance that is independent of direction. This is the same set of arguments that appear starting at Eq (7) from [21], as well as other places.

**Figure 1.**
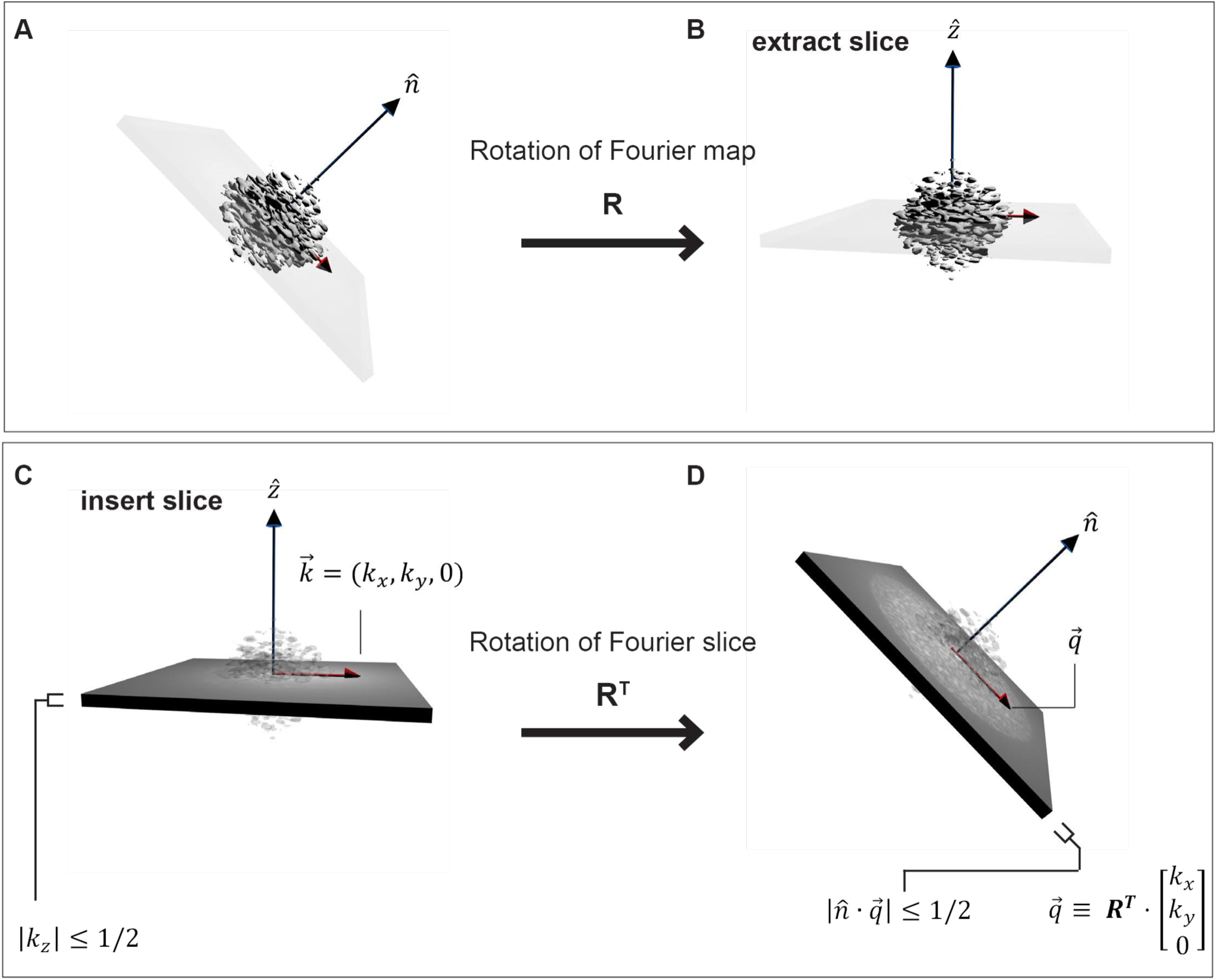
Geometry of projections in Fourier space. (A) A 3D object in its Fourier space representation is rotated by ***R***, and (B) a slice is extracted from the 3D Fourier transform (FT). Based on the Fourier slice theorem, selecting a 2D slice out of a 3D FT is equivalent to orthogonally projecting the original real-space map along the new 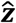 axis. (C) The data in a projection is contained in a slab of Fourier space of unit height. When considering what the data in a 2D projection, 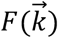, corresponds to in 3D, it is easiest to consider the mapping ***R***^***T***^, as shown from (C) to (D). Now, the coordinates on the 3D FT as shown in D are clear: 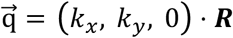. The slab condition on the projection, |*k*_*z*_| < 1/2 readily translates into the condition 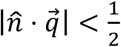, where 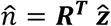.

The 3D rotation, 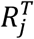, that appears in Eq (2.1) is the mapping from the 3D version, 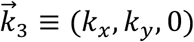 of the 2D point, 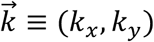 to the 3D point, 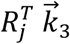, on the map, *X*, which is being reconstructed (Figure 1). The “Euler angles” for the projection, *j*, are the angles that appear in the conventional *ZYZ* representation of the rotation *R*_*j*_. The factor *M*_*j*_ (k), has been extensively described (Sorzano [21], Penczek [19]) and should be an oscillating sinusoidal function (CTF) with a frequency-dependent attenuation caused by various envelope effects. Eq. (2.1) is the generally accepted starting point for cryo-EM data.

We next redefine 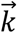 to represent points on the 3D grid, and we shift our attention to the reconstruction of the map in 3D. In the reasoning of direct Fourier reconstruction, we can form the average over the samples that are used to reconstruct each 3D grid point 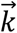 to arrive at an estimate of the 3D data point after reconstruction within the map 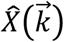:

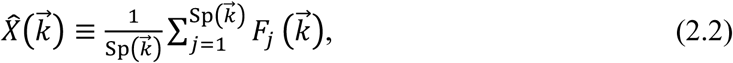

where 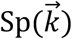 is the number of times that the particular point (in 3D) 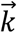 has been measured (by means of projections as described above). In a conventional direct Fourier reconstruction, both the running estimate of the reconstruction and the total weights that have been used for interpolation (that is, 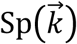) are kept as projection data is added.

One key observation is that, after substituting (2.1) into (2.2), the resulting noise is always down by a factor of one over the square root of the amount of sampling (see [21]):

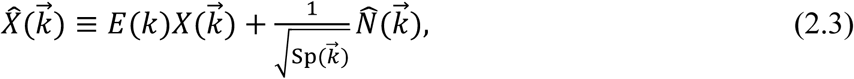

where the “renormalized” noise, 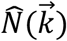, has mean zero and the same variance as the average of the variances of the constituent noise variables 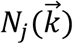. Eq. (2.3) has been written so that the variance of 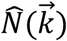 does not depend on the sampling. This is parallel to the argument which appears in [21] at about Eq. (11). We have introduced *E(k)*, which is an effective envelope and the average over the samples of the microscope influences (*M*_*j*_(k)) as well as misalignment effects. Strictly speaking, Eq (2.3) can only be approximate, but it is consistent with other approximate analyses [22].

In the typical evaluation of cryo-EM resolution, two independent reconstructions are performed to arrive at half maps which we can write in Fourier space as:

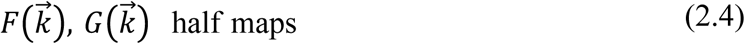

We are interested in the FSC of half maps drawn from the same statistical ensemble as in (2.3). Therefore, we consider two maps assembled as in (2.3) and then we calculate the FSC. Each half map, *F, G* should therefore be of the form given as in Eq. (2.3):

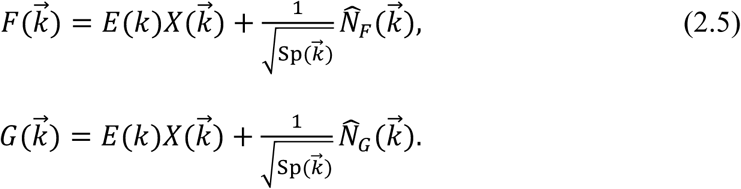

The normal prescription is to introduce the correlation of these half maps at a discrete set of wavevector magnitudes, and then examine the functional behavior of this scalar correlation as a function of this wave-vector:

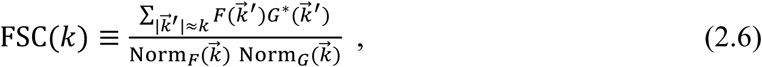

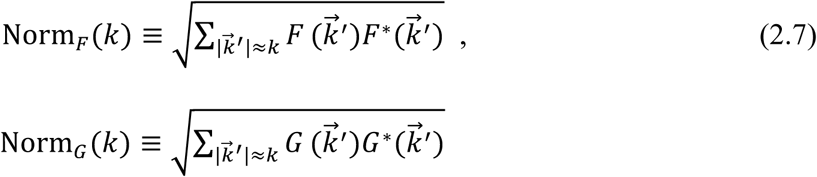

Since *F* and *G* are assumed to be statistically similar, we can write (2.6) in short hand as

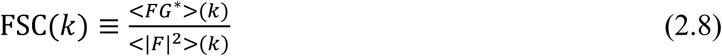

Where we have used <·> to mean the angular averages, (and equating angular and ensemble averages). Very crudely, it is the cross correlation divided by the self-correlation. For more rigor, see Sorzano et al [21], or Penczek. [19, 23].

Starting from (2.8), we can perform the familiar sort of calculation [19, 21, 24]

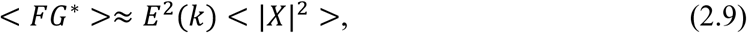

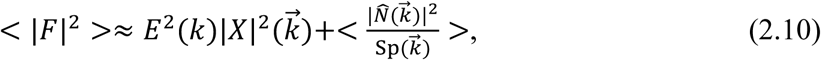

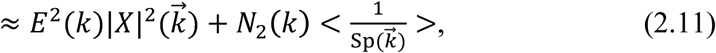

where:

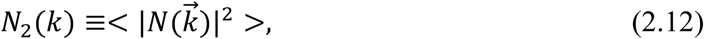

and we have decoupled the noise variance from the sampling in going from Eqs (2.10) to (2.11). There is no a priori reason to anticipate that the noise variances are related to the Euler angle assignments, so the decoupling implicit in going from (2.10) to (2.11) is consistent with standard assumptions. For the half maps, this leads to the following approximate estimate for the FSC using the above (2.8), (2.9), (2.11):

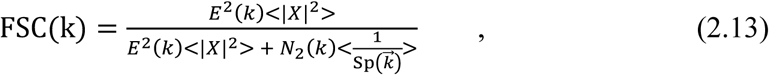

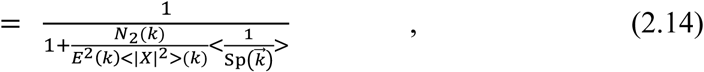

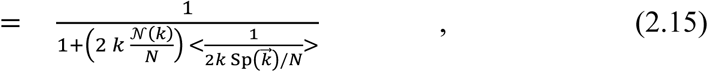

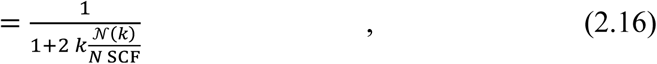

Where, in going from (2.14) to (2.15), we have defined

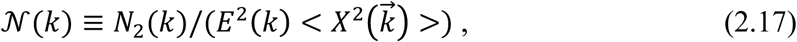

which is a noise-to-signal power ratio. In going from (2.15) to (2.16), we have defined:

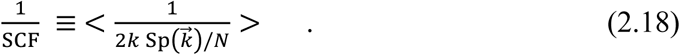

The expression (2.18) is the same as (1.2), after identifying 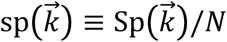, the per-particle version of the sampling. The expression for 𝒩 (*k*) is the effective noise-to-signal ratio. Notably, all the effects of sampling anisotropy are gathered into a single term: the SCF as given by (2.18).

Following previous formulations, we can define the spectral signal-to-noise ratio (SSNR):

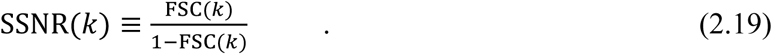

Substituting (2.16) into (2.19), we arrive at Eq (1.1):

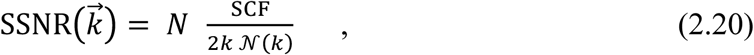

where *N* is the number of particles, SCF is the geometrical factor, *k* is spatial frequency and 𝒩(*k*) is the effective noise to signal power (given by (2.18)), whose inverse would act like the predominant component of the envelope. The reason for the regrouping of the factor 2k into the SCF expression is that the SCF is then unity in a continuum calculation for the average density of sampling for distributions that are uniform, as we will show in section 3. Eq (2.20) is essentially the same expression that appears in [24] except for the appearance of the SCF term.

Under certain circumstances, the reconstructed volume may have regions of Fourier space that have not been sampled. Two typical causes for this are: 1). The set of projection views are not well distributed (such as top views), such that Fourier voxels, even very near the Fourier origin, have not been filled. 2). The set of projection views are reasonably well distributed, but as one moves further from the Fourier origin, there are lattice sites that are not sampled. Because Fourier voxels not receiving information during the reconstruction procedure are traditionally left as zeros, there will be voxels that do not contribute to the angular averages of (2.9) – (2.11). A careful recalculation shows that the amended formula for the SSNR, as defined by (2.19), should still be (2.20), except that the angular average involved with (2.18) for the evaluation of SCF should only take place over non-zero voxels:

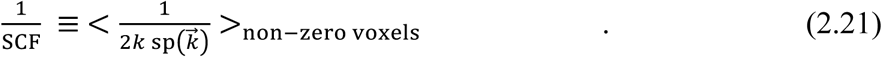

Eq (2.20) along with (2.21) are our Eq. (1.1). As an aside, we show later that 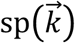 goes like 1/2*k*, so that the total sampling follows *N*/2*k*. Therefore, in the standard cryo-EM experiment, the total sampling typically will not thin to zero, and the only zeros are the result of deficient projection distributions.

### 2.2. An adjusted formula for SSNR for half maps with unmeasured data

The SSNR, based on half maps, has a drawback when some of the Fourier voxels have been left unmeasured. The voxels in each half map are typically set to zero, which leads to smooth, but artefactual maps, and may yield artificially high resolution measures. We see this in detail late in Section 5, when we look at reconstructions that are performed from projections in a 45° cone, and a percentage of randomly distributed extra projections is decreased in the sequence 10%, 3%,1%, 0%. There is a sudden increase in the improperly defined FSC resolution measure at 0%. We will defer discussion of the reconstructed data to that time. Here, we seek an adjusted SSNR expression, which allows variance to be assigned to regions of Fourier space that have been unmeasured. Consider the simplest situation, as in Figure 2, where we have represented some shell of Fourier space by *P* measured values having mean *T*, and variance per voxel, var_N_, given by the reciprocal sampling at each measured point. There are also *Q* unmeasured voxels, that are assigned 0 values in Figure 2A, and contribute neither to the signal nor the variance. Then, the ratio of signal to variance is shown in the figure: 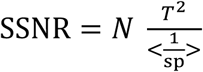 where the average is taken of the reciprocal per-particle sampling over the measured values. However, if one assigns a variance of 1 to the *Q* unmeasured voxels, and repeats the same calculation, one arrives at 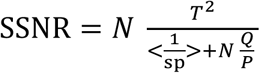. In particular, when N is large, the behavior is completely different: in Figure 2A the SSNR increases without bound, and in Figure 2B the SSNR plateaus to a finite value and is proportional to the area of measured to unmeasured region.

**Figure 2.**
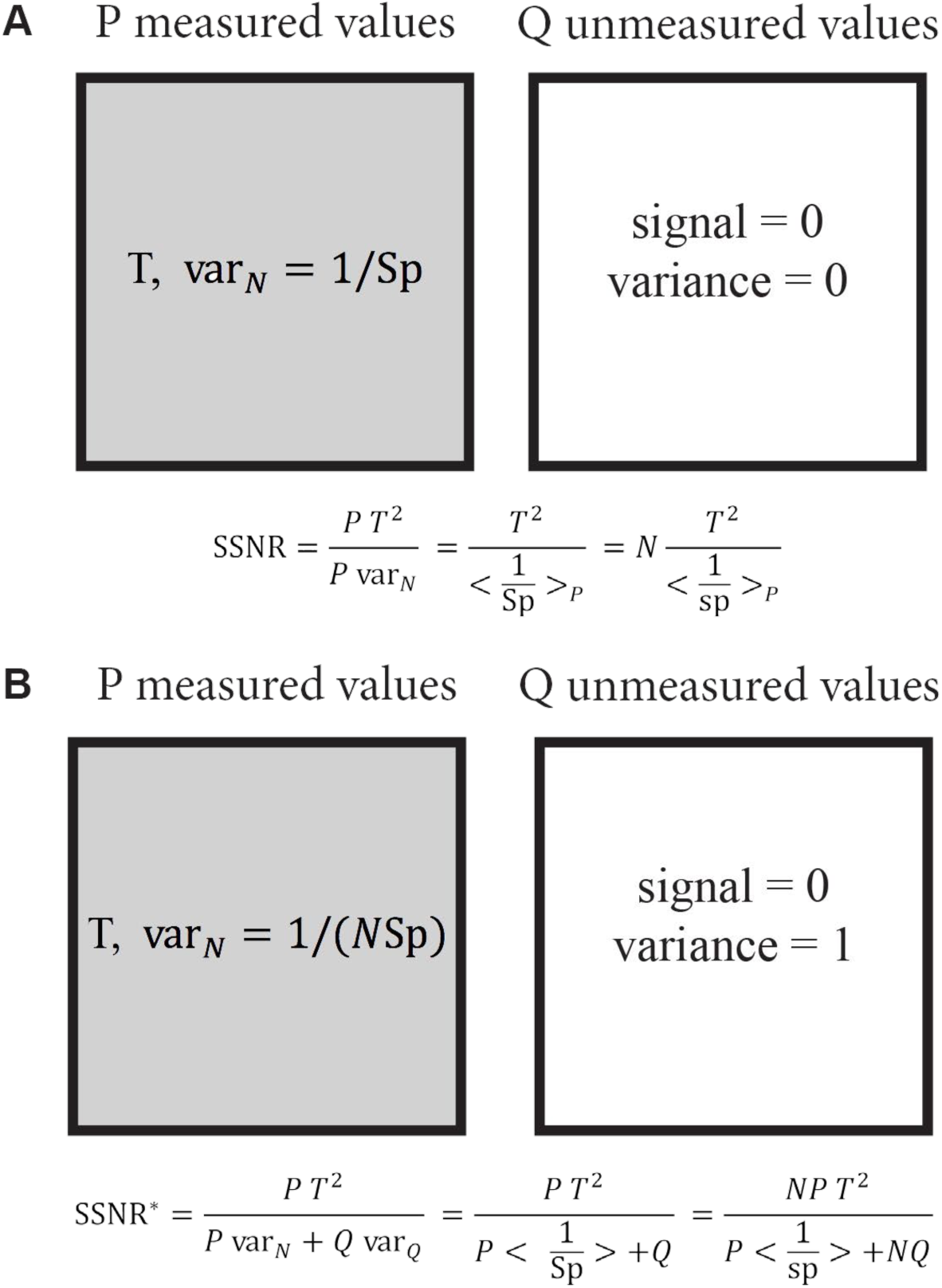
Schematic of the proposed adjustment to the SSNR formula for cases with unmeasured regions of Fourier space. The shaded areas represent the *P* values on a Fourier sphere that have been measured for a target value, T, in the measured region. The variance in the measured region is down-weighted by the total sampling, Sp, which is N times the per-particle sampling, sp. Meanwhile, the unshaded region represents *Q* unmeasured values. In (A) the unmeasured voxels are assigned zero variance, whereas in (B) the voxels are assigned variance consistent with the already measured voxels, resulting in two different expressions for SSNR. The expression in (A) limits to arbitrarily large values as the number of particles increases, whereas in (B) the expression saturates.

Generalizing the scenario in Figure 2, we consider a Fourier shell at Fourier radius *k*, and let *P*_*k*_ be the number of voxels that have non-zero sampling and let *Q*_*k*_ be the number of voxels with missing data. The total number of voxels therefore is then *P*_*k*_ + *Q*_*k*_. We calculate the adjusted values of the quantities in (2.9) and (2.10), assuming that the data with missing voxels should be allowed to have variance. Then:

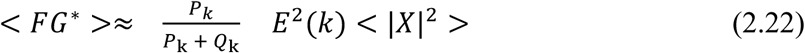

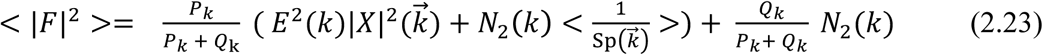

Where *E* is defined through (2.3), *N*_2_ is the noise variance defined as in (2.12), and *X*(*k*) is the target structure. Our approach for the missing data is now clear: missing voxels take on a single unit of noise unattenuated by any sampling. The fairest assignment for such voxels is one unit of variance and zero units of signal. The adjusted formula, FSC*, for the FSC then becomes:

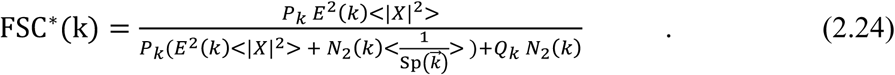

This leads to an adjusted SSNR, which we develop by starting with its reciprocal:

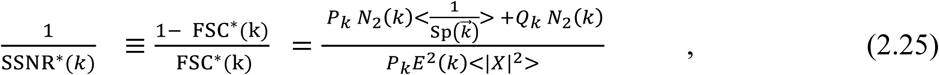

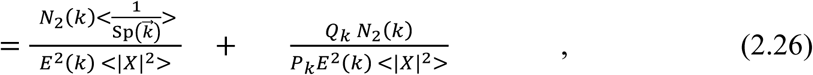

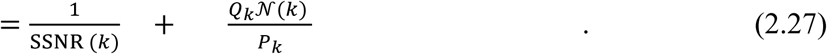

Thus:

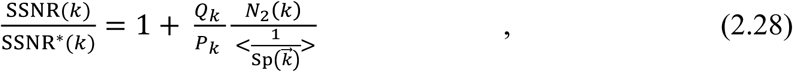

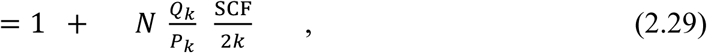

which may be rewritten:

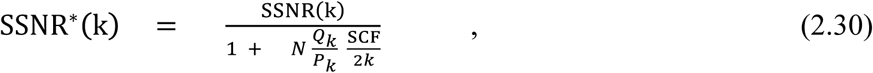

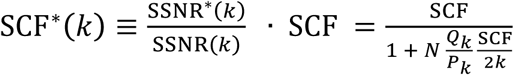

where SCF* is the expression to use in the adjusted version of Eq. (1.1): 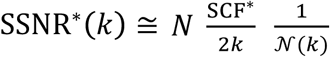. Eq. (2.30) gives an expression for reevaluating the SSNR for half maps when the original half maps have missing data. One way to think of Eq (2.30) is it shows how the conventionally constructed SSNR is inflated due to not assigning any variance to missing data. Eq. (2.30) also yields a condition by which a correction is necessitated. The ratio of occupied to unoccupied voxels at some Fourier wavevector is typically only a weak function of Fourier magnitude. This means it is also a geometrical parameter, similar to the SCF. Therefore, when

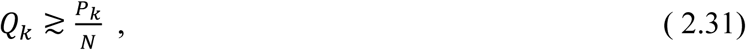

then one should have to correct with the factor in the denominator of Eq. (2.30), to obtain a more realistic value of the SSNR. The condition (2.31) is the condition that the unmeasured variance is similar in magnitude with the measured variance, which is sampled in proportion to the number of particles. Another way to write it is that we must make an adjustment when 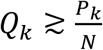. If there is a sufficiently narrow gap, then we can ignore the adjustment. In practice, if there a sampling geometry that produces true missing gaps, then any number of particles should necessitate the alternate formula.

If N is sufficiently large, then what limits the resolution is solely the gap. Adding more particles will not improve the SSNR, because additional particles will not better resolve the missing voxels, and the already measured region is sufficiently well resolved. The expression for the adjusted SSNR is most readily read off from (2.27), when the unadjusted value becomes large. Then the first term on the right-hand side can be neglected, and the reciprocal of the remaining terms taken to find the limit of large particle numbers, but with missing data:

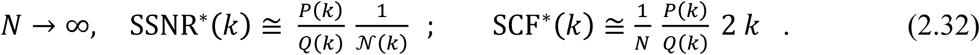

Thus, in the limit of large particle numbers, the adjusted SSNR plateaus to a value, which is the per particle envelope multiplied by the ratio of measured to unmeasured voxels. For positive k, the expression implies that the FSC* quickly drops from unity for even small 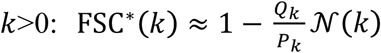 and is not improved by adding more particles. In this case, the measured voxels are perfectly well sampled, and all the variance is due to the missing values.

To summarize, we derived the relationship between the SSNR and the type of sampling distribution that is involved in the reconstruction. The latter enters the formula solely as a single geometrical factor, the SCF, given by Eqs (2.20) and (2.21) (which reiterates Eq. (1.1)), the main result of this work. In section 3, we derive analytical expressions for the SCF, and in section 5, we evaluate the efficacy of (1.1) using simulated cryo-EM datasets. In the case of missing data, we suggest an adjusted expression for the SSNR from what is usually used. This is the formula for SSNR*(k) given by Eq. (2.30).

## Section 3. Numerical and analytical forms for the sampling function, and expressions for the SCF geometrical factor

In Section 1 and 2, we showed that the entire effect of the sampling inhomogeneity on the SSNR could be incorporated into a single geometrical coefficient, the SCF. In this section, we provide numerical and analytical forms for the sampling function, as well as the geometrical SCF factor that causes decrement to SSNR curves. In section 3.1 we explain our numerical and analytical approaches for evaluating the sampling and show that they evaluate identically for appropriate cases. In section 3.2, we give continuum expressions for the sampling for several families of distributions: 1) a one parameter family of distributions with an axial symmetry, that span the complement to cones, which we term “side-like”; 2) a one parameter family of side-like distributions modulated by fluctuations in the phi angle; 3) A two parameter family of projection views that are constrained to fall within a cone of half-angle *α*, and that have, in addition, a fraction, *ϵ*, of views that are randomly scattered through the remainder of Euler space. In section 3.3, we calculate analytically the SCF for each of these distributions using the continuum formalism that we developed, which is valid when the sampling is not too small. The range of values of the SCF for “side like views” ranges from 1 (the maximum, corresponding to uniform) to 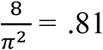 (side views). For the modulated side view cases, the SCF decreases as 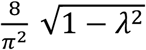, where *λ* is the magnitude of the modulation, and we restrict the modulation >1. This gives us a complete parametrization of reasonable sampling where the SCF decreases from 1 to .81 (side) to 0. For the poorly sampled top-like views, we give a closed form integral expression for SCF (*α,ϵ*) and evaluate the expression graphically. In the case when *ϵ* =0, we point out that there are typically missing values and the usual expression for the SSNR is not logical, as it neglects the variance that can be estimated for the unmeasured voxels, by using the data already measured on the same shells of Fourier space. Using the expressions that we developed in Section 2, we show theoretically that properly defined SSNR curves should always improve after increasing the sampling (by increasing the percentage of uniformly distributed views that lead to measured data in the unmeasured region). All figures of the SCF curves and dependencies on control parameters are provided accordingly.

### 3.1 Discrete and continuum approaches to the sampling function

#### 3.1.1 Discrete treatment for sampling

The projection-slice theorem [25] states that a 2D projection from a direction 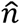 of a 3D map, is a slice out of the Fourier volume of the plane perpendicular to 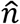 and passing through the origin, as shown in Figure 1. If we think of the map as rotated by *R* before the projection (along 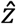), then what we term the projection direction, 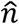, is (approximately) perpendicular to the sampled points, and is given by 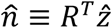. As suggested in Figure 1, each projection, 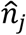, samples the set of points 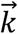 satisfying

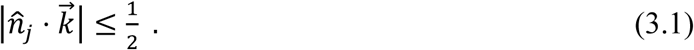

The totality of the discretely sampled points form a lattice as shown in Figure 3. Here, a single projection (in Fourier space) is taken in the 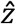 direction with Fourier magnitudes less than the real space box, *L*. Lattice sites, shown as blue dots in the *k*_*z*_ = 0 plane are considered to be sampled. Each sampled plane selects a lattice of points in this manner. Our numerical algorithm hinges on finding lattice sites that satisfy (3.1) for each projection. As we sum over projections, we increase the totality of “viewings” of each lattice site. In direct Fourier inversion, this integer number of “viewings” will correspond roughly to the reconstruction weights.

**Figure 3.**
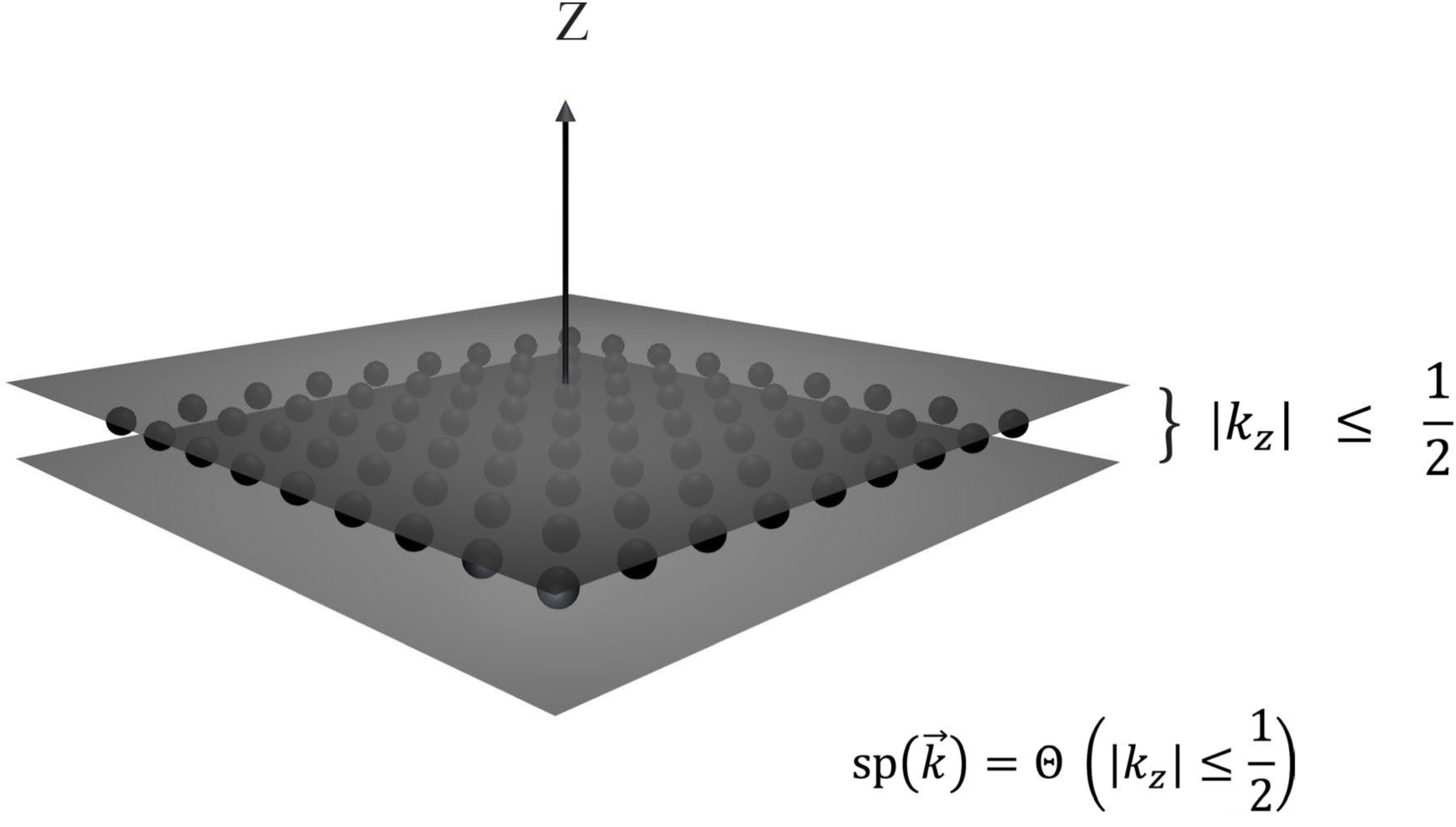
Numerical and analytical representations for sampling. A representation in Fourier space of a single projection with Fourier magnitudes less than L. Lattice sites, shown as blue dots, in the *k*_*z*_ = 0 plane are considered to be sampled. Our coarse-grained approximation is that the entire slab of unit width containing those points are sampled: 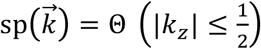, where Θ is the indicator function. Our numerical algorithm hinges on finding lattice sites that satisfy the generalization of this condition: 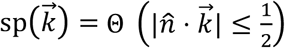 for any arbitrary projection direction given by 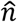. For analytical calculations, we further approximate this as a delta function 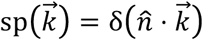, which satisfies the proper normalization.

The number of times a particular 3D point, 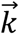 is sampled, we term 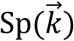, and is therefore given by the cardinality of the set of the projections, that for a given 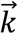, satisfy the criterion of Eq. (3.1). Therefore:

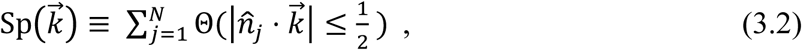

where Θ is the indicator function (see Glossary). The per-particle sampling function we define as:

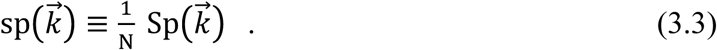

Eq. (3.1) is what is used numerically to find the sampling at each voxel, wherein the vector to each voxel is checked against every projection to see if the dot product between this vector and the unit direction given by the projection is sufficiently small (less than ½ in magnitude).

We investigate suitably many approximations that the sampling function emerges as a quantity that independently affects the SSNR (and only coupled to average microscope effects: not individual CTFs per particle, for example). It is our hypothesis that this level of approximation is sufficiently useful to enable understanding the effect of anisotropy on resolution.

#### 3.1.2 A continuum treatment for sampling

We wish to formulate the expressions analytically whenever possible. Toward this end, we recast (3.2) using Dirac delta functions, which will provide continuum calculations that are both useful and accurate. For a single projection in the z-direction, we would like to employ

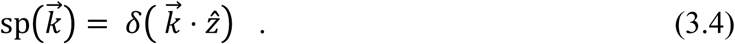

Generally the Dirac delta function is considered to be 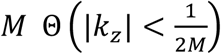, in the limit that the parameter M becomes arbitrarily large, whereas we have taken *M* as simply unity in (3.1). The delta function analytical approximation is crude, but satisfies the proper normalization.

The generalization of (3.1) for continuum calculations using the idea in (3.2) yields

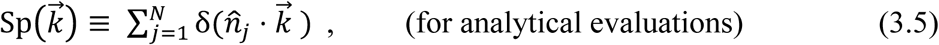

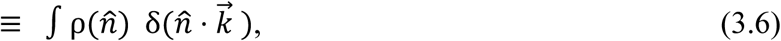

where 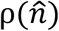 is a measure on the distributions of projections parametrized by 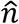: (in this case, 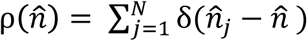, which is discrete, but generally 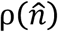 may be continuous). Eq. (3.5) is the continuum approximation when the length of the side of the box, which we will use as L, can be considered to be much larger than 1. This is sufficient for many of our analytical treatments and development of formulae, since we are often working far from the Fourier origin. In Eqs. (3.5), we consider Fourier space to be dimensionless (unitless), which is a common practice. To reintroduce units, if one has, in 1D, 200 voxels of voxel size 1 Å per side, then each Fourier space voxel will have width 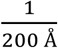 and the largest distance from the Fourier origin will be 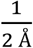 (this is the Nyquist frequency). The average sampling across a shell at fixed Fourier magnitude can be derived using our continuum treatment. Starting from Eq. 3.4:

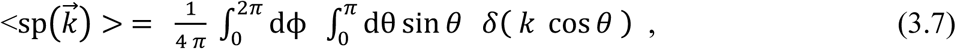

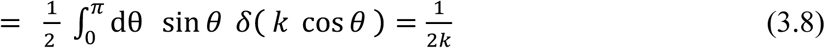

This is a natural result: placing planes (Fourier slices) into volumes, the density must fall off as one over the Fourier radius. A more thorough derivation is given in Appendix A.1, including an interpretation of the geometrical factor 2, which couples to the k dependence. Eq. (3.8) is also the sampling per particle for a uniform set of projections, but Eqs (3.7) and (3.8) hold for any distribution of projections.

#### 3.1.3 Consistency between numerical and analytical expressions for sampling

As a check of both our code and analytical implementation, we tested the total amount of sampling in our volume by placing 50,000 projections in a box of size LxLxL with L=41. We calculated the integer sum, S, over all the sampling at all the points, and evaluated *S*/4*L*^2^ numerically to be 1.19. To develop an analytical expression or this idea, we can write

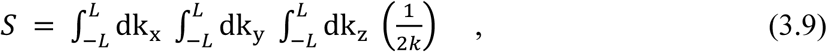

where k is spatial frequency. This is the average amount of intersection of arbitrarily oriented planes with a cube of side *2L*. In the appendix A, we show that the integral evaluates to

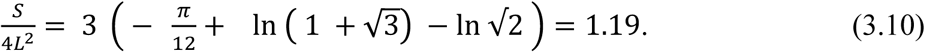

This corroborates our numerical result described above.

### 3.2: Sampling Function for three different distributions in continuum representation

We calculate the sampling function for three different distributions. The first is the case of the complement to a cone (which we term side-like). The second is for side views with a modulation in the azimuthal Euler angle. Finally, we also calculate the top-like cases, where a certain fraction of uniform views is also included. The side-like views and side-modulated views are each governed by single parameters: i) the cone half angle, *α* and ii) the modulation parameter, *λ*. The top-like family of distributions is governed by two parameters: once again, the cone half-angle, *α*, and a parameter, *ϵ*, to cover uniform projections in the complement to this region.

Figure 4 shows the projection distributions that will be described in this section, including a schematic representation of the projection distribution (top row), the population of Fourier space through slice insertion (middle row), and the experimental sampling map derived from 10,000 insertions (bottom row). These are displayed for different sampling schemes, including the uniform (Figure 4A), side-like or complement to cone (Figure 4B), side (Figure 4C), side-modulated (Figure 4D), and top-like (Figure 4E).

**Figure 4.**
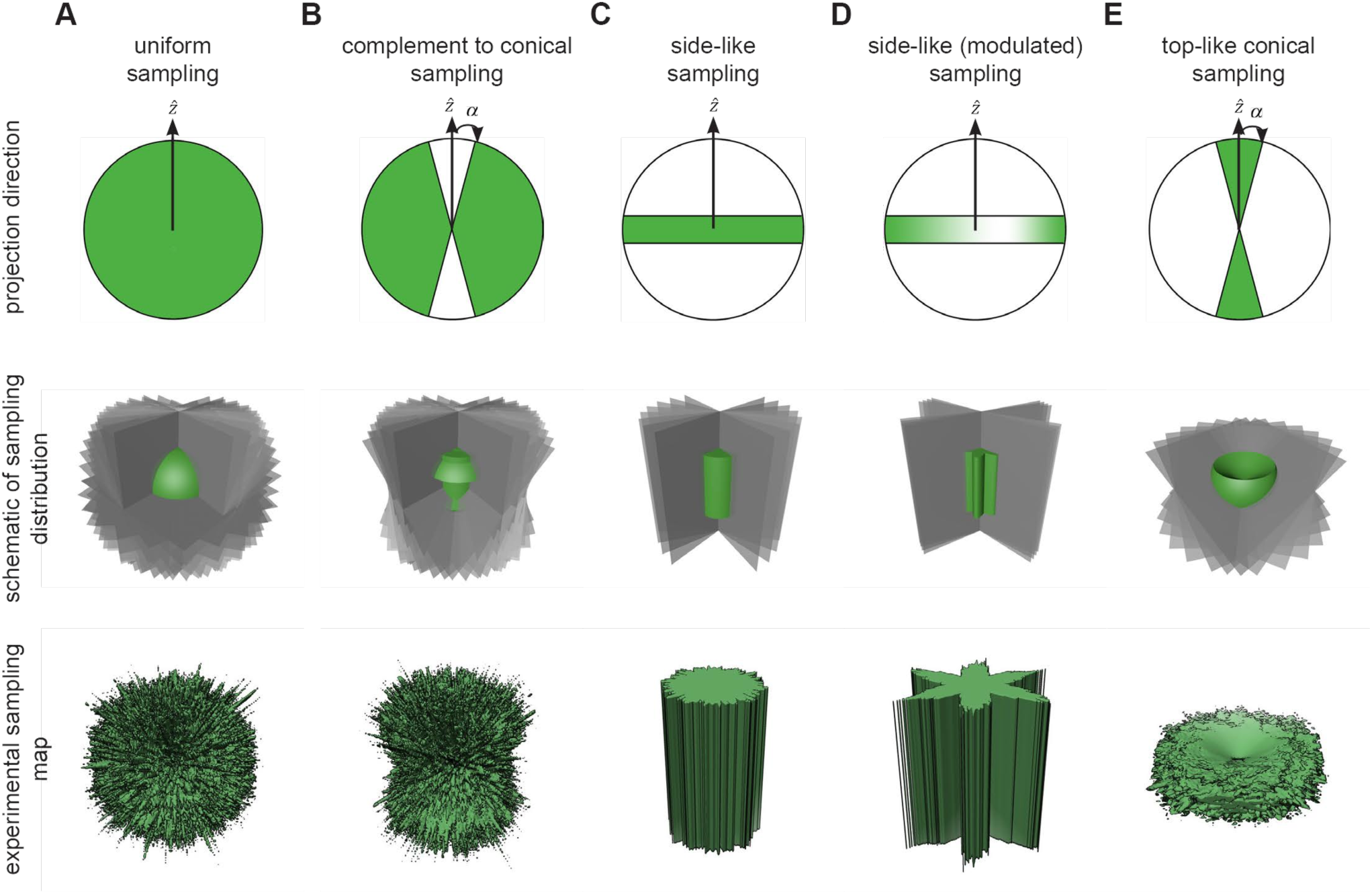
Projection distributions used to evaluate sampling. The first row denotes the directions of the projections (green); the middle row provides a schematic of the sampling (Fourier slices are in gray and the sampling map schematic is in green); the last row shows the experimental sampling map from 10,000 slices inserted into the 3D FT. (A) The uniform sampling distribution evenly covers the entirety of reciprocal space. (B) “side-like” projections are uniformly drawn from the complement to a cone of half angle, *α*. The uncovered region is orthogonal to the cone and lies along the X/Y plane. (C) Projections are drawn from the side, which corresponds to the *α* →*π*/2 limit of case B (Euler angle θ=90°). (D) In addition, we incorporated azimuthal oscillations (Euler angle ϕ is modulated), which increases the fluctuation of side view sampling. The depth of the oscillations is governed by a parameter, *λ*, and we term this scenario modulated side-views. (E) Projections are drawn from within a cone of half angle α. Unlike the other cases, the top-like distributions always have missing conical regions of Fourier space related to the size of the half-angle *α*. For this reason, we ultimately include an additional parameter, *ϵ*, which represents the fraction of projections scattered randomly over the projection sphere.

#### 3.2.1 side-like cases (*α)*

For the side-like case, we have

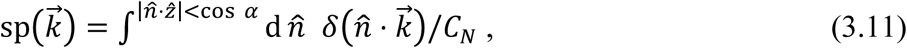

where *C*_*N*_ is a constant to ensure the normalization (3.8), leading to (see Appendix B):

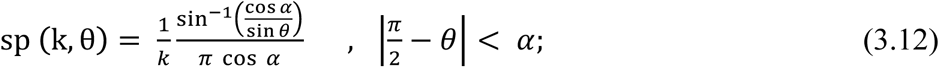

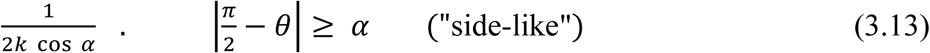

The distribution for side views can be selected by taking the 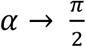 limit to arrive at:

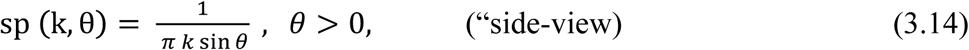

Along the z-axis (that is, *k* sin *θ*=0), sp should have the same value as at the origin, which is 1.

#### 3.2.2 modulated side-views (*λ*)

A set of modulated side view projections can be written as a density distributions:

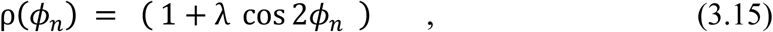

where *ϕ*_*n*_ is the azimuthal angle for a projection direction 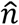. This gives rise, therefore to a sampling given by:

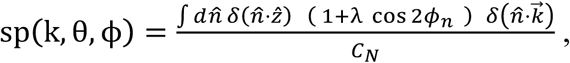

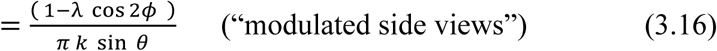

We describe in appendix A.3 how to select a set of projections with this form, using the cumulative distribution function.

#### 3.2.3 top-like cases (*α, ϵ*)

Finally, we consider sampling for the top-like cases:

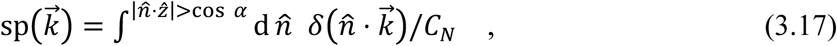

which leads to (see Appendix B):

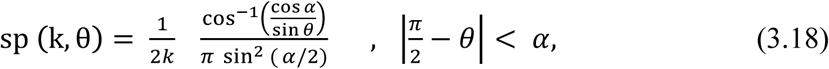

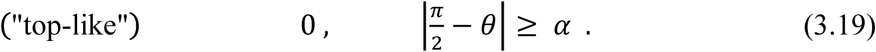

Taking *α* ≪ 1 leads to arbitrarily large values of sp: 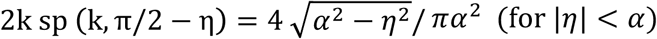. Once again, the sampling needs to be truncated to unity, when *α*=0, *θ* = π/2, that is in the xy plane, if the top-view is taken along the z-direction.

These distributions have missing data, and so we can calculate, for each shell, the ratio of filled, P, to unfilled voxels, Q.

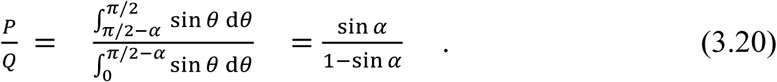

We will need Eq. (3.20), to compute SSNR*, as argued in section 2, because there is missing data and develop an adjusted formula for top-like SCF distributions in the next subsection.

We also can evaluate the top-like cases, when we add a random distribution of projections, so as to fill in the missing data. Then:

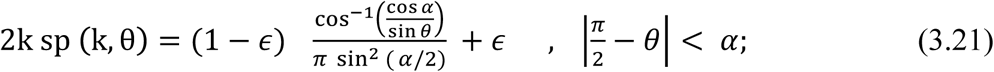

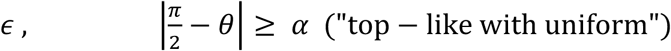

where *ϵ* is the fraction of projections that are distributed randomly, and the rest fall in the original cone of half-angle *α*.

### Section 3.3 The Sampling Compensation Factors for the three different distributions

The SCF is defined via:

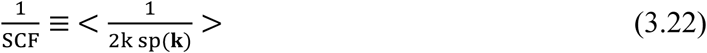

where < ·> is the average over solid angle regions that have non-zero values of the sampling. We can evaluate this numerically for the “top-like” and “top-like with uniform” distributions, and analytically for the “side-like” and “modulated side-view” cases.

In Appendix A.5, we evaluate Eq. (3.22) using (3.14) to arrive at:

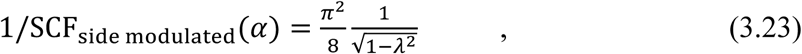

which ranges from arbitrarily large values to 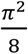 (when *λ* = 0; no modulation). In practice, the sampling never achieves a continuum of values, so the expression given by (3.23) cannot be used for *λ* very close to 1.

One can also show:

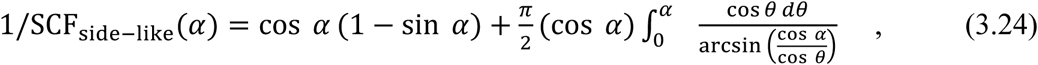

ranges from 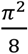 (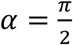 side views) to a value of 1 (*α*= 0, uniformly distributed views).

Figure 5 shows schematically the behavior of the SCF for the side-modulated and side-like cases. It is shown there how to continuously vary the SCF from its low values (corresponding to side-modulated cases with sampling suffering from deep pockets) to its highest value unity (for uniform views).

**Figure 5.**
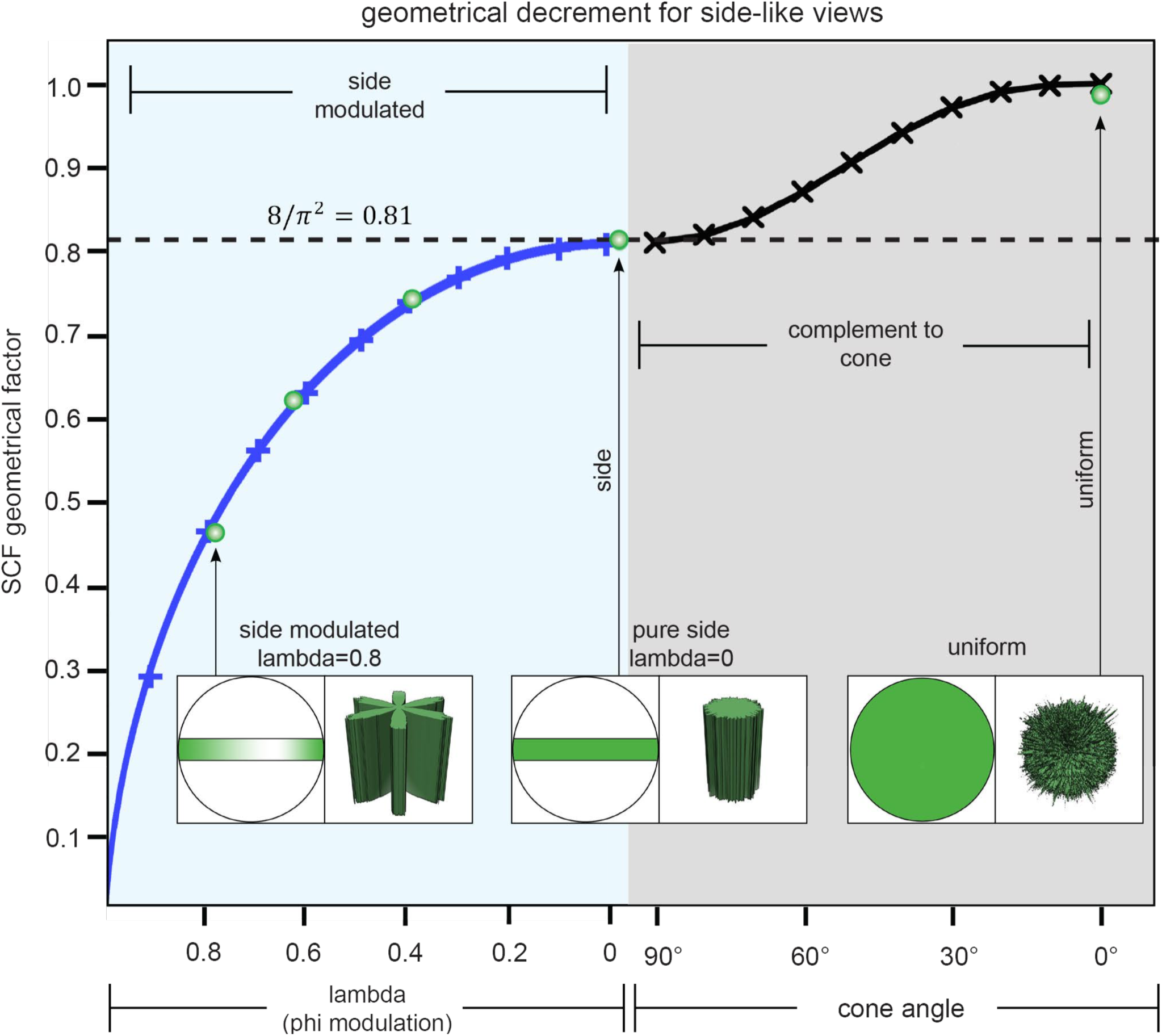
Geometrical decrement of the SCF for side-like views characterized by thorough sampling. The SCF is a single geometrical factor, which forms an approximate estimate for how the SSNR is decremented due to deficits in sampling. We plotted SCF_side modulated_(*λ*) for modulated sets of side views (see Eq (3.23)), as well as SCF_side-like_(*α*) for side-like views (see Eq (3.24)). The approximations inherent in Eq. 3.23 are no longer valid for *λ* very close to 1, and therefore the plot is not shown for *λ* < 1, SCF > 0. The projection views and sampling maps are shown in three typical cases: (i) *λ* = 0.8 side-modulated, where there is a deep pocket in the sampling, (ii) pure side views, where the contours of equal sampling are cylinders, and (iii) uniform views, where the SCF attains its maximum value of 1. Green open circles represent the numerical evaluations of the SCF, and their correlation with our continuum calculations (represented by the curves) reinforces the efficacy of both approaches.

Finally, we want to calculate the quantity that represents the top-like situation. Since much of the region is zero, the normalization from (3.22) must be carefully calculated and leads to:

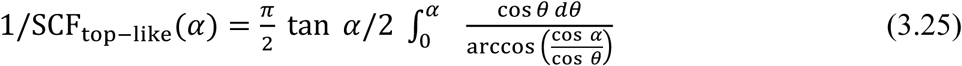

The right-hand side of (3.25) ranges from 1 (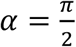, uniformly distributed views) to 0 (for *α*= 0, purely top views). The asymptotics are 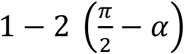 for 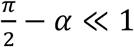and 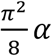 for small *α*. Thus, we have a set of analytical expressions for SCF that can run from arbitrarily small to unity and from unity to arbitrarily large levels. However, any distribution with SCF > 1, involves distributions with missing data. Ultimately, the more relevant attribute, will be SCF* which relates how the correctly adjusted SSNR* is decremented due to the sampling. Thus, the SCF* is bounded by 0 ≤ SCF* ≤1.

Repeating with the additional random projections gives a drastically different value for the SCF for the singular change of adding partially uniform perturbations, because now all (or most) of the Fourier points have at least some sampling.

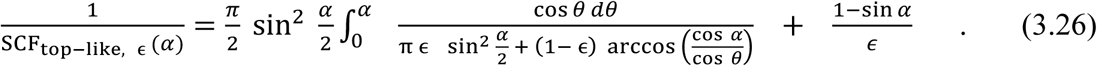

For such top-like distributions, it is interesting to compare 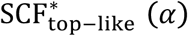 from (2.31) with SCF_top-like, ϵ_ (*α*) for small but finite ϵ from Eq. (3.26). From Eq (2.31), (3.20) and (3.25), we can derive the *ϵ*=0 quantity:

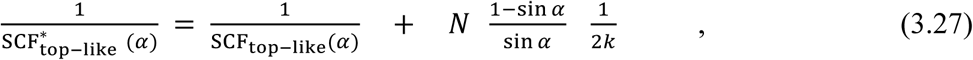

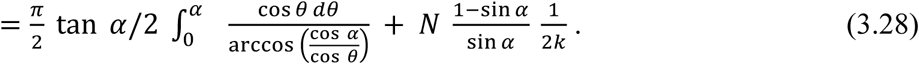

Note that 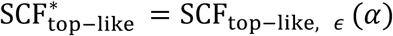, when 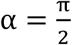. For small *ϵ*, but large *N*, the second terms of both (3.26) and (3.28) dominate and we get:

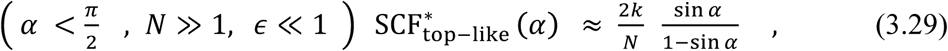

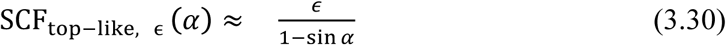

The crossover between these expressions occurs approximately when

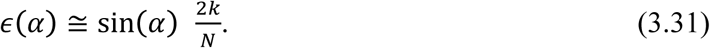

The situation for the decrement in the correctly adjusted SSNR is depicted In Figure 6, for the poorly sampled cases. The Eq. (3.28) is the lower bounding curve in gray (*ϵ*= 0). Otherwise, the curves represent Eq. (3.26). There is no crossover, unless epsilon is sufficiently small: in the figure there are only three crossings of the curves. For *k* = 15, and N = 10^4^, Eq. (3.31) implies 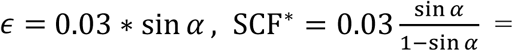 is the crossover between curves.

**Figure 6.**
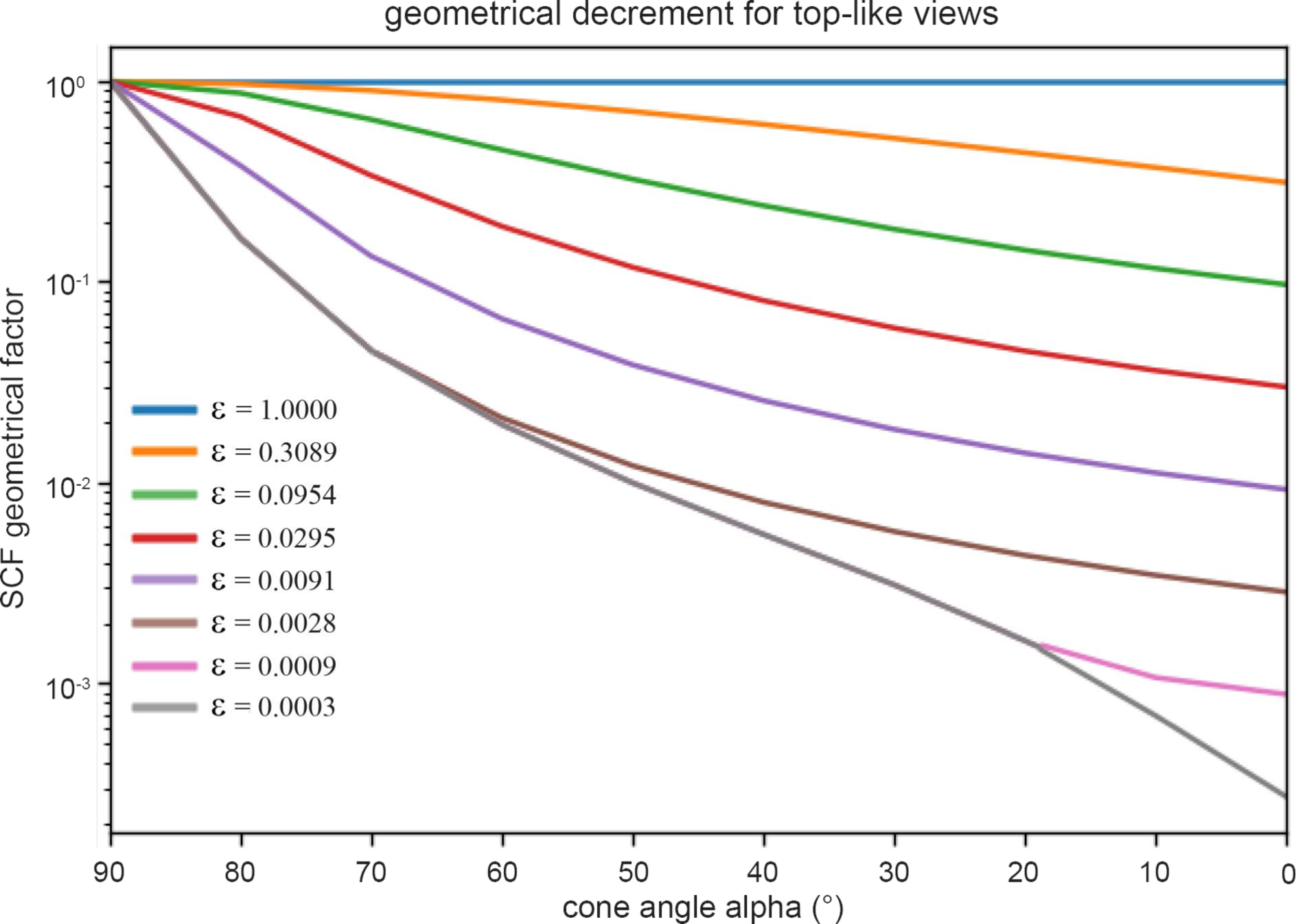
Geometrical decrement of the SCF for top-like views characterized by poor sampling. Plot shows the predicted SCF as a function of cone size and fraction of randomly sampled projections. The attenuation of SCF due to conical sampling is typically more severe than for well-sampled cases. The angle, *α*, represents the cone (in degrees) that predominantly contains the projections, except for a fraction, *ϵ*, which represent the percentage of projections that are distributed uniformly over the projection sphere. Typical distributions are shown later in Figure 8. The curve that bounds the whole set from below, is given by Eq. (3.28), but is otherwise given by Eq. (3.26): the plotted SCF^*^ is the maximum of the two different expressions. This crossover between expressions is given at the bounding curve (gray) when 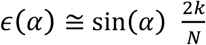.

The last expressions tell the entire story of missing data. If data is missing in some sizeable region, the adjusted SSNR is drastically reduced. However, even a small fraction of random perturbations starts to quickly reintroduce signal. If there is a gap, the SCF is increased by a factor

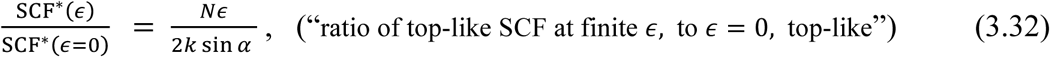

by adding back a fraction ϵ worth of random perturbations. For *N* = 10^4^, *k* = 15, α = 45°, *ϵ* = 0.01, the RHS becomes 4.7, which is a huge jump over such a small change in ϵ. Conversely, having an empty region of Fourier space gives much lower SCF* than a lightly sampled Fourier space.

## Section 4. Relationship between SSNR and the number of particles N in a reconstruction

In Section 2, we derived the relationship (2.20) (or equivalently (1.1)), which is the estimate of the SSNR in terms of the sampling. There are two aspects of the latter: the cumulative extrinsic effect due to the number of particles in the data, and the shape of the distribution of the sampling (or projection directions), an intrinsic quality. When Fourier space is reasonably sampled everywhere, we can assign a single parameter to each of the extrinsic and intrinsic qualities of the sampling: *N*, the number of particles, and SCF, the sampling compensation factor, defined as in Eq (1.2). The SSNR is seen to be proportional to each quantity, with the SCF attaining its maximum value of unity when the distributions of projections are uniform.

In this section, we revisit the dependence of the SSNR on *N*, the number of particles, when every other aspect of the problem is held constant:

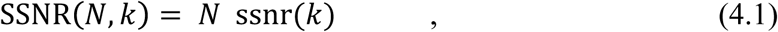

for some function, ssnr, which is the form of Eq. (1.1) with 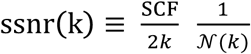. Eq. (4.1) is the familiar way that the signal in a noisy system should accrue, if *N* represents the total number of measurements. The per-particle SSNR, which depends on many factors that corrupt the final reconstruction, is observed to be quite rigid and independent of *N*, as previously noted [20]. The resulting universal curve, ssnr(*k*), includes multiple components inherent within the cryo-EM pipeline that attenuate resolution: attenuation due to the microscope transfer function, detector noise, incorrect image orientation assignment, structural heterogeneity, among others. The consequence is that the number of particles needed to obtain a higher resolution using the same collection scheme can be determined from a single SSNR curve, provided that the curve is sufficiently smooth at the desired resolution: indeed, smoothness of the SSNR curve might be another possible criterion for resolution. The universality of the SSNR/*N* curves is akin to the familiar Reslog [26] or Guinier [20] analyses.

### 4.1 Linear dependence of SSNR on N

According to Eq. (4.1), dividing the SSNR by the number of particles results in a universal per-particle curve. To test this idea, we looked at sequences of FSC, equivalently SSNR, curves for reconstructions using successively larger number of particles, N, for data from an experimental dataset contributing to a 2.9 Å reconstruction of the eukaryotic large ribosomal subunit [27]. Figure 7A shows a total of seventeen experimental FSC curves, from N=7000 to 70000 particles. The series of FSC curves collapse to a universal curve via SSNR/N, as predicted by 4.1, where 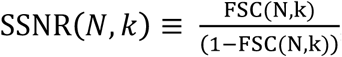, as shown in Figure 7B. Although this idea has appeared formally in many places [20, 26, 28, 29], we have not noted the explicit construction of such universal curves, as highlighted here. For smaller values of particle number, *N*, the ssnr(*k*) curve loses continuity at smaller values of resolution and limits our ability to calculate the necessary number of particles to achieve higher resolutions, as described below.

**Figure 7.**
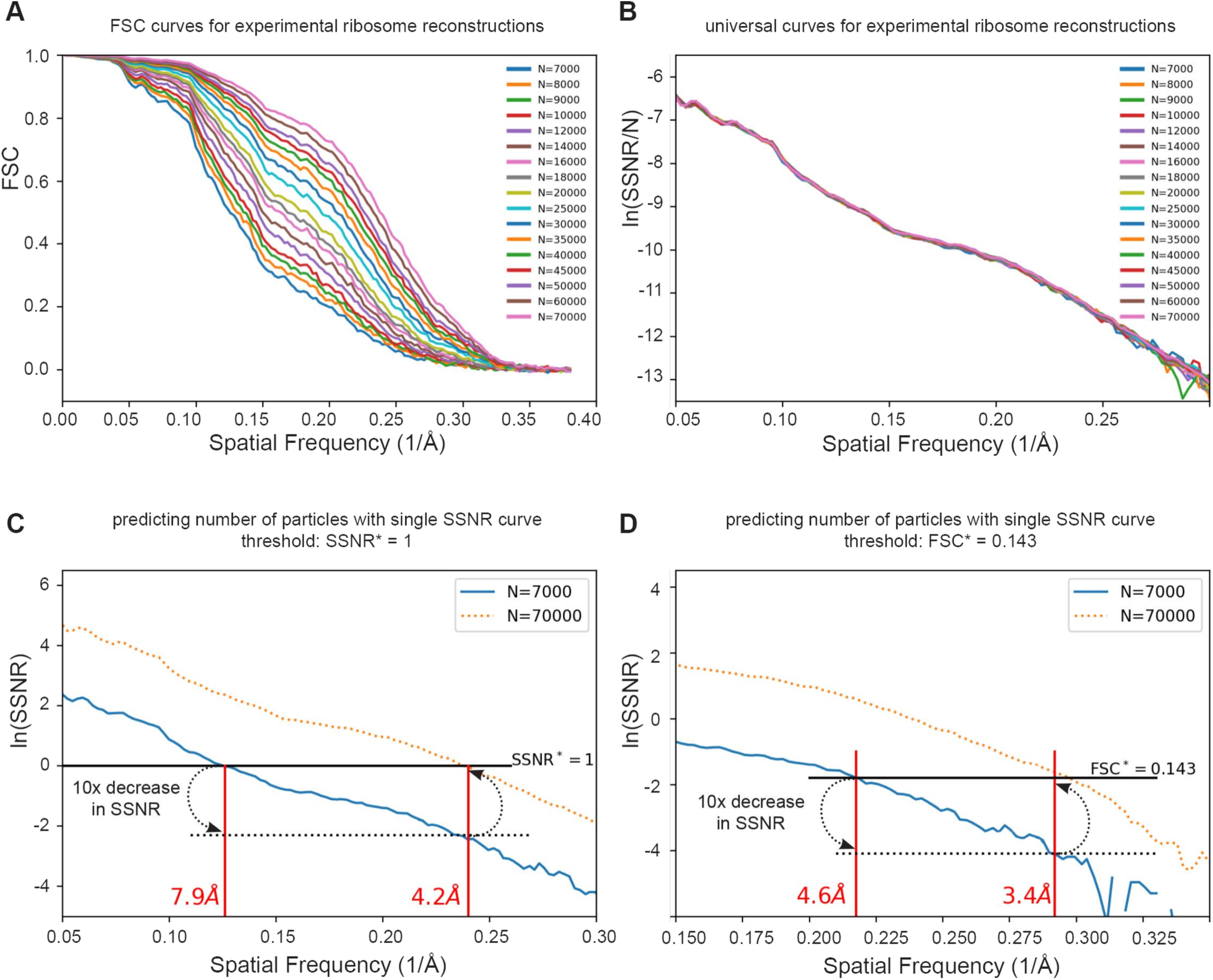
Predicting the number of particles based on per-particle SSNR curves. (A) We reexamined 17 of the FSC curves with the largest numbers of particles from Passos et al [27]. (B) A plot of the per-particle ln(SSNR/N) shows that these 17 distinct curves collapsed approximately to a single curve. (C-D) Since each curve contains essentially the same information, we can estimate the number of particles needed to achieve a target resolution. For example, one may wish to know the approximate experimental resolution by increasing the number of particles by tenfold from 7,000 (solid blue line) to 70,000. (C) using the SSNR*=1 threshold (solid black line), one would find the intercept corresponding to a 10x decrease in ln(SSNR) (dotted black line), and plot that back onto the solid SSNR* = 1 line. The experimental ln(SSNR) curve for 70,000 particles (dotted orange line) shows a correspondence between the prediction and the experimental intercept. (D) The same argument approximately holds for the FSC* = 0.143 criterion, or for other thresholds.

### 4.2 Number of Particles Necessary for Reconstruction

Eq. 4.1 can be used to predict the number of particles necessary to attain a given resolution for a general envelope function, ssnr (*k*), derived from a single SSNR curve. A common scenario that is encountered during cryo-EM data collection is one in which the experimentalist asks whether the current approach is conducive toward achieving a target resolution, given a fixed amount of collection time. Our claim is that, there is some N_0_ so that for N = N_0_, we can construct the curve ssnr(k)= SSNR(N_0_, k)*/N*_0_ and arrive at a reasonable estimate predicting the necessary number of particles (it is conceivable to make a lower estimate for the necessary N_0_, but this is beyond the scope of the current discussion). Thus, for a resolution criterion, FSC = FSC*, one arrives at an implied criterion, SSNR = SSNR* = FSC*/(1 – FSC*) (If FSC* = 0.143, or 0.5 then SSNR* = 0.167 or 1.0 respectively). Next, one defines *k*_T_ to be the target resolution. Then the necessary number of particles, *N*_*T*_, to achieve the target resolution is given by:

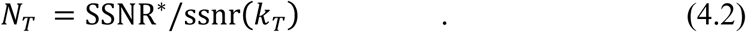

Graphically, we can make a construction on a semilog plot of the original SSNR curve, and realize that

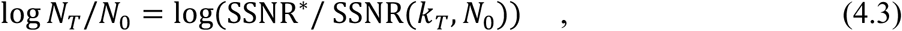

which follows from Eq. (4.1), which implies *N*_*T*_/*N*_0_ = SSNR(*N*_*T*_, *k*_*T*_) / SSNR(*N*_0_, *k*_*T*_), and SSNR (*k*_*T*_, *N*_*T*_)= SSNR*. Now, Eq. (4.3) can be used to graphically find the number of particles needed to achieve a target resolution, since the shift from the current resolution to a target resolution gives the ratio in the number of particles to increase to. Unlike other methods discussed in this section, this makes no assumptions about the functional form of the per-particle SSNR, ssnr(*k*): it can be exponential (Reslog) or Gaussian (Guinier) or indeed hold to any shape.

The idea is demonstrated for the ribosome sequence of reconstructions in Figure 7C. For convenience and in line with standard assumptions in the cryo-EM literature, we used the same two FSC criteria described above of 0.5 and 0.143, which is equivalent to an SSNR condition of SSNR^*^ = 1 and 0.167, respectively, and analyzed the SSNR curve corresponding to 7000 particles. Using SSNR^*^ = 1 (or equivalently FSC=0.5), the resolution is measured to be 7.9 Å. To obtain the necessary ratio of number of particles required for reaching the target resolution of 4.2 Å, and using this same criterion, we can measure the difference on the log plot, which is 2.3 or log(10), that is one decade. Therefore, the prediction is that 10 times the original number of particles are necessary to obtain a reconstruction at 4.2 Å. When the orange dotted curve that corresponds to 10x particles is then inspected on the plot, the prediction is corroborated, since the resolution of the 70K particle FSC curve, where the orange dotted curve intercepts the SSNR^*^ condition, matches to the predicted 4.2 Å. The identical analysis is repeated using the FSC=0.143 criterion in Figure 7D.

Finally, we note that the SSNR is inversely proportional to the geometrical SCF factor, so that distributions with lower SCF (more fluctuations in the sampling) require larger numbers of particles. Under typical data collection procedures, the SCF is fixed by the sample preparation and microscope conditions, and one cannot easily consider the use of the SCF as an independent control parameter that can be conveniently varied. The exception would be to tilt the specimen, which would alter the orientation distribution, and thus the SCF [16].

### 4.3 Comparison of graphical methods (Guinier, Reslog, and per-particle SSNR curve)

The Guinier [20] and Reslog [26] formulations are popular for extrapolating the number of particles necessary for reconstruction. We would like to understand the relationship between these graphical constructions and the per-particle SSNR curves. We give a thorough analysis of the Guinier analysis, and see that the Guinier assumptions essentially also imply (4.1), but restrict ssnr (*k*) to a Gaussian form. Our method is seen to be slightly more general, but in typical usage, identical to these, based on the argument below.

The prescription in Guinier analysis [20] is to estimate the number of particles needed to achieve a given resolution, and mark this on a semilog plot of N as a function of the square of the spatial frequency, and repeat. This procedure is presumed to form a line, which can be extrapolated to find the number of particles to achieve a desired resolution. That is, knowing *N*_1_, define *k*_1_ implicitly by SSNR (*N*_1_, *k*_1_) = SSNR^*^, where SSNR^*^ is the fixed value of SSNR that demarcates resolution as described above, and define *k*_2_ similarly. The Guinier assumption is that:

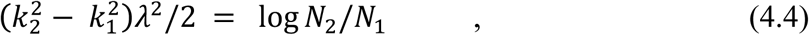

for some constant *λ*, for resolutions of interest corresponding to *k*_1_, *k*_2_. That is, along the fixed contours of SSNR, the change in the square of the resolution is proportional to the logarithm of the ratio of the number of particles used to achieve the SSNR criterion. By means of such a construction, one can estimate the number of particles needed to achieve a higher resolution. Eq (4.4) is easily solved formally as

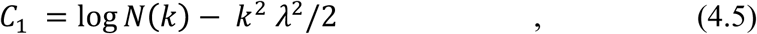

where *C*_1_ acts like a constant of integration, which depends on the SSNR, which is held fixed in the construction. This implies (exponentiate 4.5):

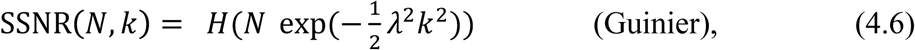

for some function *H*. Every set of SSNR curves of the form (4.6) will yield (4.4). The only reasonable choice for *H* is linear, which matches our result (4.1), when ssnr(*k*) from (4.1) is a Gaussian. We should point out that in light scattering, Guinier plots are used as a low frequency approximation where the various physical parameters can rigorously be argued to hold to the damped Gaussian format indicated by (4.6) [30].

The Reslog analysis [26] is very similar and leads to:

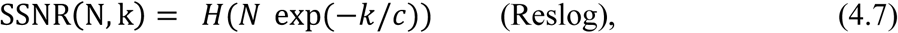

for some constant wavevector *c*. Once again, the only reasonable choice is a linear function, *H*, leading to Eq. (4.1) with an exponential form for ssnr(k) in Eq. 4.1.

Heymann [24] made an identical argument to arrive at our Eq. (4.1) and used the Guinier analysis, based partly on the formal results on blurring [31] and other envelopes [22]. As an aside, much like multiple time scales [32] can create an effective 1/frequency noise in physical systems rich with multiple time scales, with so many differing sources of noise in cryo-EM, it may be that, depending on the experimental circumstances, the linear behavior is equally valid to the quadratic behavior for governing the log of the envelope. In any case, Heymann suggests the Gaussian form for ssnr above: 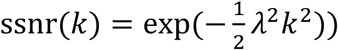, consistent with the Guinier analysis. Although Heymann arrives at Eq. (4.1), he does not arrive at our Eq. (1.1), because the possible anisotropy is not discussed, and therefore he uses the expression for the uniform distribution of the per-particle sampling (1/2*k*) which is our Eq. (3.8) and Eq. (9) of Heymann [24].

## Section 5. Decrement of SSNR through non-uniform sampling

In Section 4, we observed that the SSNR depends in two ways with the sampling; the extrinsic part governed by the number of particles *N* (as already has been discussed in the literature) as well as the type of the sampling governed by the geometrical factor of the sampling map, which we have termed the SCF. In this section, we test whether the SCF (or SCF^*^) has the predicted effect on the SSNR as described by Eq. (1.1) and explained in section 3.3. We look at sequences of reconstructions of two proteins that vary in their size and shape: the influenza hemagglutinin trimer and human apoferritin, for all the situations for which we calculated the SCF (or SCF*) values in Section 3.3. In each case, we compare the SSNR curves of reconstructions versus the baseline case, which is a set of uniformly distributed views.

### 5.1 Methods

#### 5.1.1 Generation of projection distributions

We generated a set of 10,000 projection Euler angles for sequences of different sampling distributions, each of which is described in section 3.2. We evaluated three different schemes for modulating the projection distribution and comparing to the uniform distribution, as depicted in Figure 8. For the well-sampled side-like sequence, we used pure side views and modulated side views with a set of modulation parameters given by *λ* = 0.4, 0.6, 0.8, and 1.0, (Figure 8A). For the first of the more poorly-sampled cases, we selected top-like projections, distributed within varying cone sizes of half angular width (5°, 30° and 45°), and fixed a small amount of random projections (3%) distributed evenly across the rest of Euler space (Figure 8B). This scenario evaluates the effect of increasing cone size. For the second of the more poorly-sampled cases, we fixed the cone size to be 45° and added random assignments of 0%, 1%, 3% and 10% evenly distributed projections across the rest of Euler space (Figure 8C). This scenario evaluates the effect of increasing the amount of random projection “sprinkling” in the presence of an otherwise fixed distribution.

**Figure 8.**
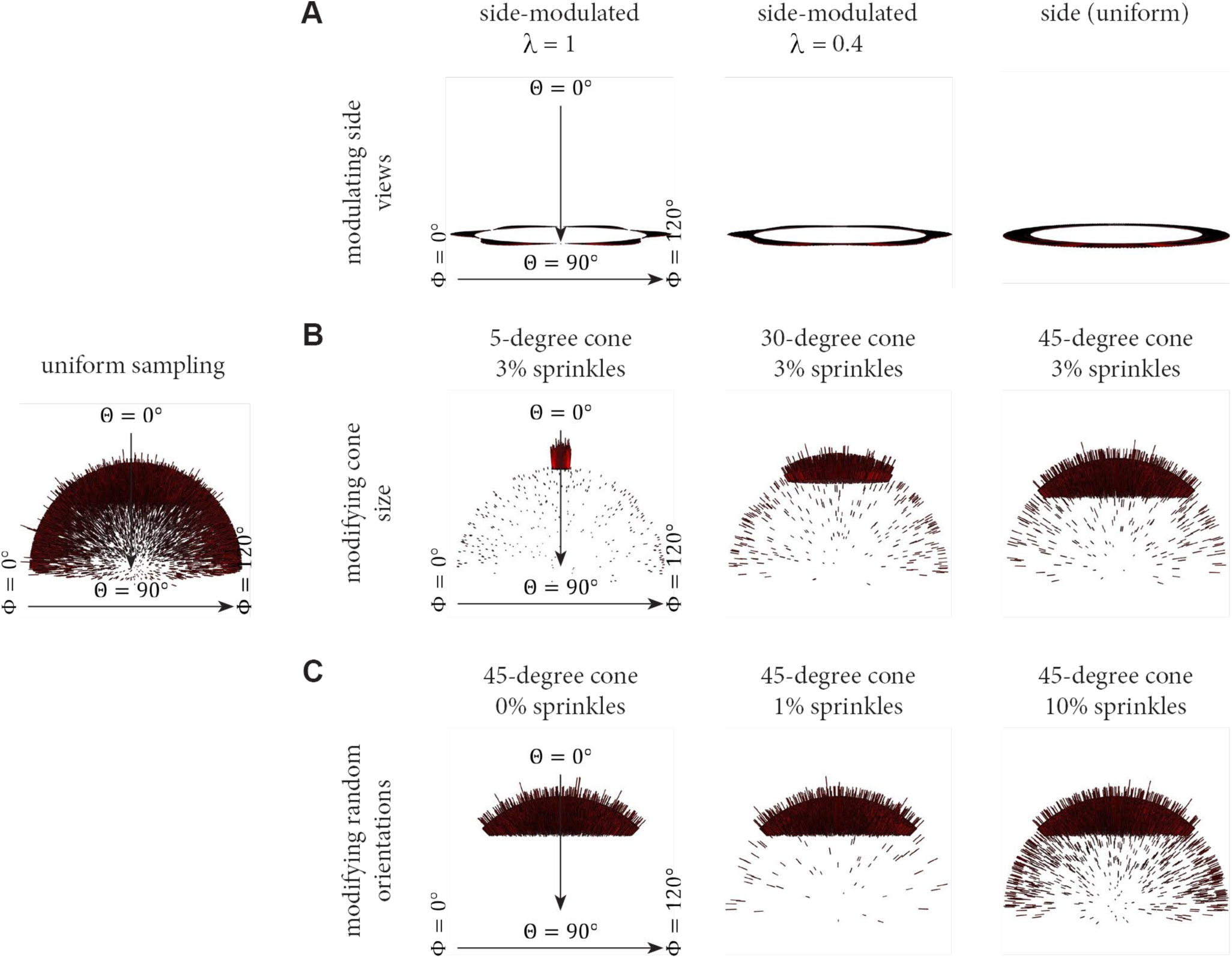
Euler angle sampling schemes used in this study. The projection distributions described in Figure 4 can be modulated to vary specific parameter and evaluate distinct conditions. For each scenario, Euler angle distributions for 10,000 projections are displayed. A uniform distribution of views across Euler space is shown at left for comparison. (A) The side-modulated case, whereby the Euler angle θ=90°, but ϕ is modulated in accordance with the modulation parameter, *λ*. This scenario corresponds to the transition between Figure 4C-D. (B) The top-like case, whereby the size of the cone is varied, and there is a fixed amount of randomly distributed views across the rest of Euler space. (C) The top-like scenario, whereby the size of the cone remains constant at 45°, but the amount of randomly distributed views is varied across Euler space. The experimental results corresponding to these three cases will be described in Figures 9-11.

#### 5.1.2 Synthetic data generation with distinct projection distributions

To test our idea relating the effect of a single geometrical parameter and the SSNR, we generated synthetic datasets corresponding to two proteins of varying size and shape, namely the hemagglutinin (HA) trimer and apoferritin. The synthetic data generation followed previously described protocols [16, 33-35]. Briefly, 10K projections were generated from cryo-EM maps of either HA or apoferritin, according to the viewing directions that were described in Section 5.1 above. These projections were shifted and rotated at random and noise was added. Next a distribution of CTFs were applied to the 2D projections, followed by an additional layer of noise to arrive at an SNR approximately equal to 0.05. This SNR is consistent with experimental cryo-EM data [36]. A reconstruction was performed with the known orientations using the Frealign software, and the usual FSC was calculated between half maps. In parallel, the angle assignments were used to calculate sampling maps, as described in Section 3 (and shown graphically in Figure 4). From the sampling maps, the SCFs were calculated numerically by implementing (3.22).

### 5.2 Results comparing decrements predicted by sampling and reconstructions

We tested how well the SCF geometrical parameter, based solely on the projection directions, could predict the decrement of the SSNR, with all other aspects of the problem held constant.

#### 5.2.1 Side-like and side-modulated sampling cases

We first proceeded to test the predictive ability of the SCF on well-sampled cases, where most of the values of the sampling remain reasonably high and each index point is sufficiently sampled, e.g. above 20. From a theoretical perspective, we should expect that the ideas set forth are most accurate in the this scenario. This is a typical case in cryo-EM reconstructions, even if some views are dominant. In the well sampled cases, all the structure factors remain at play, so we expect that the formulae relating SSNR to SCF are reasonably accurate. The situation is presented in Figure 9, where we describe the effect on reconstructions for a uniform case, for side views, and for modulated side-views. For both reconstructions of HA and apoferritin, in comparison to uniform, the SSNR curves are attenuated for side sampling in accordance with the amount of sampling inhomogeneity (Figure 9A-C). Side views have a range of sampling values over the surface of a Fourier sphere of radius, *k*, from the on-axis values to those on the orthogonal plane with a max-min ratio of *πk*. For the modulated side-view case, the ratio is even larger: *πk*/(1 − *λ*), where *λ* is the strength of the modulation. Nevertheless, the agreement between the decrement in SSNR and the SCF, as shown in the table in Figure 9D, is acceptable for both HA and Apoferritin.

**Figure 9.**
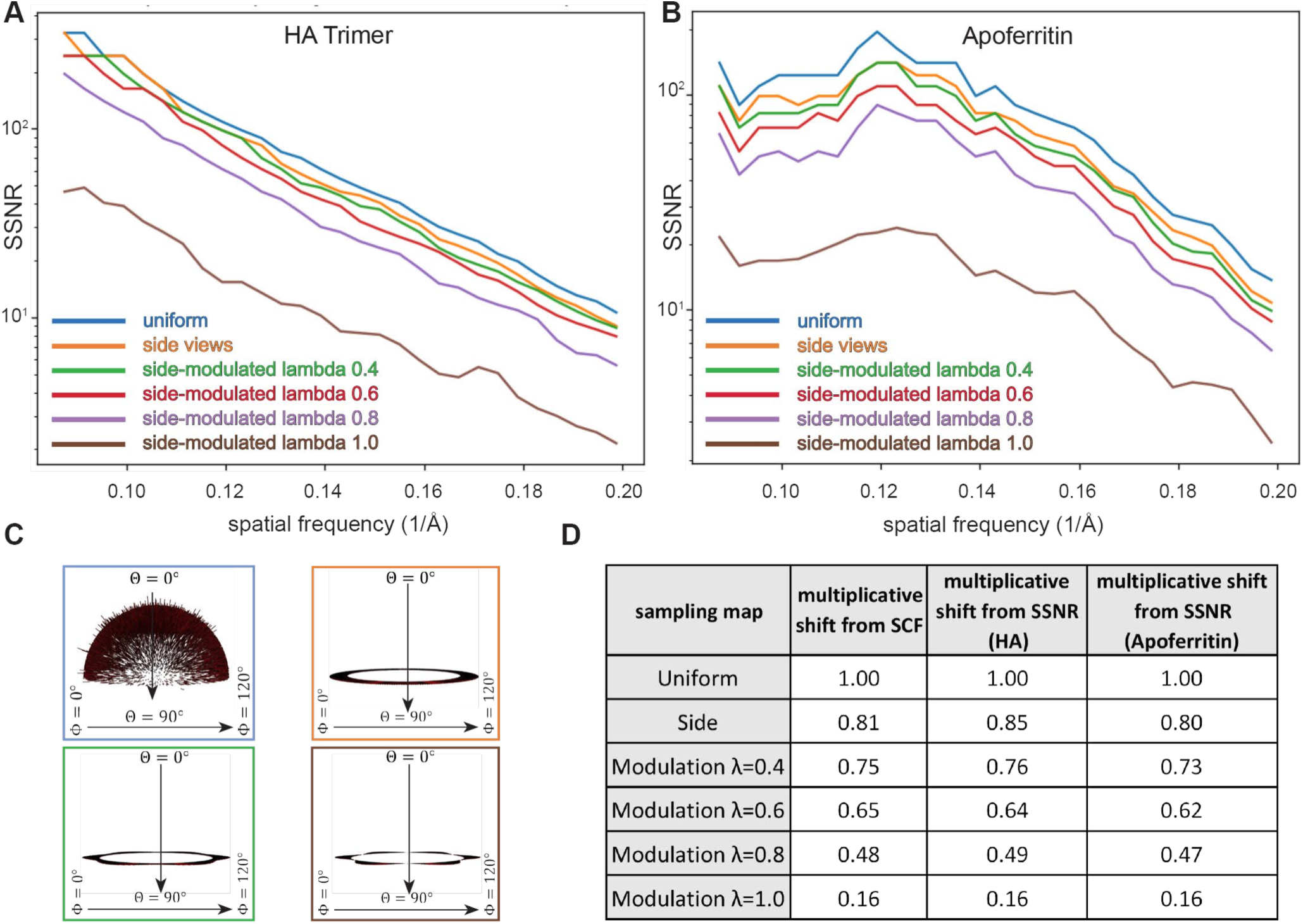
Attenuation of the SSNR due to the projection distribution for side-like cases characterized by thorough sampling. (**A-B**) Two synthetic datasets corresponding to (**A**) the HA trimer and (B) Apoferritin were reconstructed, as described in section 5. The decrement, which is reflected by the shift in the SSNR due to the different type of sampling distributions is displayed. (**C**) Euler angle distribution profiles corresponding to selected cases from A-B. (**D**) Table showing the decrement in SCF and the multiplicative shift from SSNR. For each of the 6 different sampling distributions, the numerical and analytical forms for the SCF agree (except for *λ* = 1, where we only have a numerical formula), and thus only a single multiplicative shift from SCF is indicated.

#### 5.2.2 Top-like sampling cases for varying cone sizes

We then proceeded to describe cases that would reflective of a predominant top view, and for this reason constructed the two-parameter family of distributions described by *α* and *ϵ*, where the former represents the half-angle of the cone from which the projections are drawn, and the latter represents the percentage of uniform projections besides those drawn from the cone. First, we vary the size of the cone, while fixing 3% uniform sampling across the rest of Euler space. Figure 10A-C shows how the SSNR is attenuated for reconstructions generated from such top-like distributions containing a fixed amount of sprinkled projections. In these cases, the maximum to minimum sampling can be so large as 1 + 4/(*πϵα*), for small *ϵ, α* according to the analytical formula. Nevertheless, in Figure 10D, the multiplicative shift determined from SCF (both numerically and from formulae) approximately matches the decrement in SSNR.

**Figure 10.**
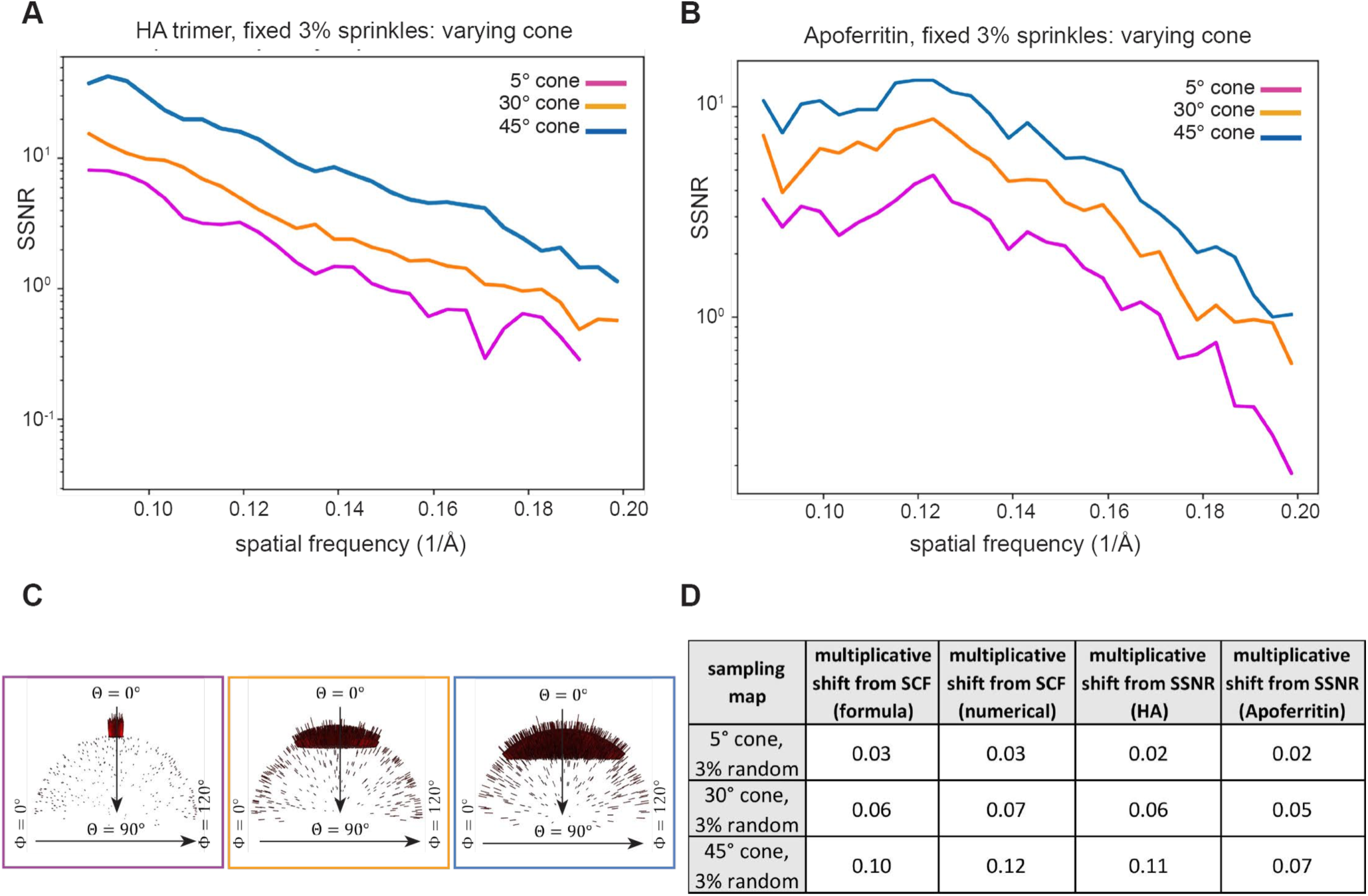
Attenuation of the SSNR due to the projection distribution for top-like cases characterized by varying cone sizes. As in Figure 9, for both HA and Apoferritin, we examined the effect of the SCF on the SSNR for poorly sampled cases, where the projection distributions were constrained to varying cone sizes, but the number of random projections was fixed at 3%. This leads to Fourier space being sparsely sampled around the z-axis in each case. (**A-B**) Decrement of the SSNR for (**A**) HA and (**B**) with (**C**) the corresponding Euler angle distribution profiles. Even though there is a low percentage of views in certain regions, the total sampling is not small, because the total number of particles is 10^4^. (**D**) Table showing the decrement in SCF. In each case, there is a crude agreement between the analytical expectation for the SCF, the numerically calculated SCF, and the shifts of the SSNR.

#### 5.2.3 Top-like sampling cases for fixed cone size and varying fraction of randomly sprinkled projections

Finally, we took the same two parameter family as in Section 5.2.2, but examined a fixed cone size, and varied the fraction of random projections. Figure 11A-C shows how the SSNR is attenuated for reconstructions generated from such top-like distributions containing a fixed cone and varied number of random projections. The first observation from this data, as we explained in Section 3, is the artefactual increase in the SSNR for cases with completely missing data (black dotted curve in Figure 11A-B). This stems from the singularity in the theory for how the SSNR is typically defined, and a separate formula is needed to properly account for the variation that is implicitly missing, in half maps created from sets of projections with missing data. The adjusted formula from Eq. (3.27) pushes the SSNR curve to the appropriate ordering of the curves, where increasing the sampling always increases the SSNR (black solid curve in Figure 11A-B). Theoretically, the other curves (1%, 3%, 10%) should not be adjusted, because there is sufficient sampling to add information to the missing regions. In practice, there is also a small shift in those curves, which is not shown for the sake of clarity. The second observation from this data is that, for cases with large gaps in Fourier space, a small amount of additional projections goes a long way in increasing the SSNR. This is not surprising. Even in the early days of reconstructions, it was realized that, for under-sampled cases, adding small amounts of information to deficient parts of Fourier space greatly improves the ability to solve the reconstruction problem [37]. The experimental attenuations of the SSNR are also in line with the geometrical decrement of the SCF in continuum calculations (compare Figures 11 and 6). As in the previous cases described above, Figure 11D shows that the multiplicative shift determined from SCF (both numerically and from formulae) approximately matches the decrement in SSNR.

**Figure 11.**
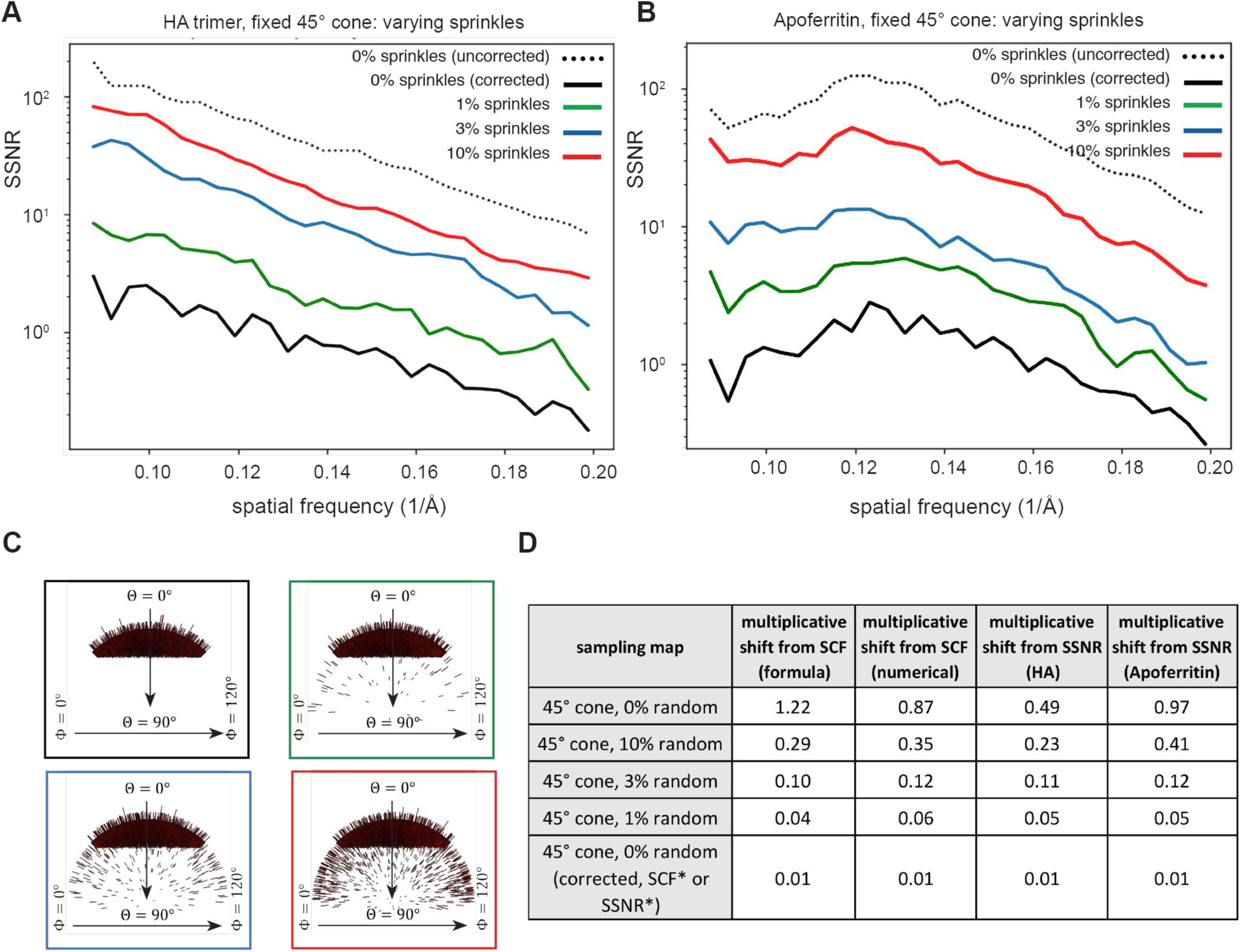
Attenuation of the SSNR due to the projection distribution for top-like cases characterized by varying random views. As in Figures 9, for both HA and Apoferritin, we examined the effect of the SCF on the SSNR for poorly sampled cases, where the projection distributions were constrained to the fixed cone size of 45°, but the percent of random projections varies from 0%, 1%, 3% to 10%. (**A-B**) Decrement of the SSNR for (**A**) HA and (**B**) Apoferritin with (**C**) the corresponding Euler angle distribution profiles. (**D**) Table showing the decrement in SCF. For the first four rows in the table (and the corresponding SSNR curves in A-B), the SCF, as defined theoretically by Eq (3.25) for 0%, and Eq (3.26) for the 1%, 3% and 10% cases, and numerically by Eq (3.22), approximately describes the observed change in the SSNR, as given by the last two columns. There is the one serious issue, as discussed in the text, that the SSNR with completely empty regions of Fourier space is significantly higher for 0% uniform rather than 1%. The text explains a logical correction, given by the SCF theoretically by (3.27) and numerically by the same algorithm. The correction to the SSNR is shown in the last row and appropriately predicts a large multiplicative shift from uniform, as would be expected for such a poorly sampled case.

## Discussion

In this work, we show that non-uniformity of the set of projection views drives down properly defined global resolution measures. Our calculations are based on standard assumptions, that there is some envelope that seems to stabilize for values less than 10 Angstroms [20]. The SSNR resolution measure estimates the ratio of the signal power to the signal variance. Using ordinary statistics, we expect that the variance per voxel will be decremented by the sampling. Therefore, if we assume that the noise variance approximately decouples from the sampling, then the average over Fourier shells of the reciprocal sampling arises naturally in the expression for the SSNR, leading to Eqs (1.1) to (1.2). Thus, the measure for the efficacy of sampling that we advocate, the SCF, emerges naturally, if we wish to isolate the effects of the geometry of the sampling on the resolution. The incorporation of the SCF is the step that distinguishes our calculations from similar calculations, such as [24].

A typical cryo-EM reconstruction procedure carries along information that can be represented by three maps: two half-maps and a sampling map that can be created from knowledge of the angle-assignments or that can be taken to be the map of reconstruction weights in a direct Fourier reconstruction. From these maps, one can estimate up to second moments and continue to combine information to arrive at more refined reconstructions. Ultimately, one arrives at a mean map, variance map, and sampling map, or three pieces of information per voxel. If there is missing data, then there is a pathology in the way that SSNR is typically defined. Although defining the mean of the missing values to zero is acceptable (and forms the best estimate of the original structure), setting the variance to zero is illogical, since there is enough information to give a better estimate. We find a self-consistent correction to the ordinary SSNR and showed in section 5 that the redefined SSNR always increases with more uniform sampling, as should be expected.

We also demonstrated the linear dependence of the SSNR on the total sampling, which is governed by the number of particles. This was implicit in earlier analyses of Guinier or Reslog, as shown in the mathematical description of section 4, but takes on a simpler form here. We show that these latter constructions imply a definite functional form for the SSNR, which is more restrictive than necessary. Indeed, we provide the mathematical argument, that one can estimate the number of particles necessary to achieve a higher resolution, using the same collection strategy, but with a single SSNR curve, provided that the curve is sufficiently continuous over the resolution ranges in question. This has value during data acquisition, since it can inform the experimentalist how a given collection might be altered or abandoned based on the goals of the experiment, and the prediction is achieved without the need to recalculate reconstructions using particle subsets.

There are several major implications from the current work. Most importantly, the direct relationship between sampling and global resolution in single-particle cryo-EM implies that any deviation from uniformity *always* drives down the SSNR, and thereby leads to an increase in the number of particles that are required for attaining a specified resolution. There is a persistent problem of preferred specimen orientation (and consequently non-uniform projection distributions) that appears to affect the vast majority of single-particle reconstructions [6]. This means that virtually all data sets are characterized by non-ideal imaging and image processing conditions. As dictated by Eq. 1.1 (also Eq. 2.20) the experimental situation therefore requires optimizing two parameters – the experimental “envelope” *as well as* the sampling distribution. Here, we use the term “envelope” in a broad sense to encompass all of the factors that attenuate experimental resolution. These include, but are not limited to, beam coherence, ice thickness (and its effect on the background signal-to-noise ratio), quantum efficiency of the detector, residual specimen movement that is not corrected by motion correction, errors in computational orientation assignment, structural heterogeneity, and any other factors that generally attenuate experimental resolution, as measured by the FSC. In addition to the envelope, the sampling distribution matters. To reach the hypothetical resolution limit for small particles [38], it is therefore essential to not only improve hardware and software, but also techniques for specimen preparation, in order to maximize sampling uniformity on cryo-EM grids. Some effort toward this goal is ongoing [7], but more needs to be done. Along these lines, the more symmetric the particle, the more orientations are sampled during the reconstruction process. Therefore, symmetry does not merely multiply the number of particles in the data in accordance with the symmetry group; the improvement in sampling for symmetric particles also contributes to gains in SSNR by virtue of improvements to the SCF. Thus, symmetry has a dual effect in improving both data quantity and quality. In part for this reason, cases like AAV [2] and Apoferritin [3] have pushed the resolution limits and are associated with very low temperatures factors (or slowly decreasing envelopes) in the data.

Beyond attenuation of global resolution, the extent to which the map suffers as a consequence of incomplete sampling is currently unclear. Specimens with high C-or D-fold symmetry that are characterized by pure side views are, strictly speaking, anisotropic. However, the effect at the level of the reconstructed map is negligible, and the experimentalist should not notice differences in structural details if one were to directly compare to a map reconstructed from a uniform sampling distribution. Nonetheless, as we show in figure 9 and emphasize throughout this work, the SSNR for pure side views is still attenuated in comparison to uniform by ∼20%, and thus the amount of data required for reaching certain resolutions is increased by approximately the same percentage. Beyond the simple cases, there are multiple factors that currently complicate an exhaustive analysis of experimental maps characterized by different symmetries and sampling geometries. First, it will be necessary to decouple the effect of anisotropy (in its strict definition, impacting directional resolution) from the attenuating effect on *global* resolution. More worryingly however, we believe that there may, in certain extreme cases of missing data, be systematic bias in the reported resolution in the field, caused by artefactual inflation in the FSC (for example, as observed in figure 11). In part for this reason, we introduced the FSC* and SSNR* criteria, which compensate for missing views in Euler space and report a more realistic value of resolution for the pathological cases. FSC* and SSNR* can, in principle, be extended to highly under-sampled orientations that may be prevalent in experimental situations. Implementation of these criteria to experimental data, and a careful analysis of the underlying sources and resulting statistics, will be the subject of future work.

Experimental improvements to the sampling distribution can be achieved by tilting the specimen inside of the electron microscope. However, this comes at a cost of degradation in image quality [16]. The direct relationship between sampling and resolution indicates that any attenuation due to sampling can now be compared with other types of experimental attenuations, for example due to beam-induced movement, ice thickness, errors in the image processing pipeline, etc. Thus, a natural direction will be to quantify the resolution gains caused by improvements in orientation sampling, as compared to resolution losses caused by degradation of image quality during tilted data acquisition. Such studies will help to quantitatively establish an optimal tilt angle for any dataset containing a given sampling distribution.

Finally, The SCF provides a direct means by which to evaluate a sampling distributions, with an intuitive scale ranging from 0 to 1. We propose the use of the SCF for evaluating Euler angle assignments for sets of particles that produce 3D reconstructions in cryo-EM.

## Acknowledgements

PRB would like to thank Pawel Penczek for conceptual explanations pertaining to the current work. PRB and DL would like to thank David De Rosier for discussions. PRB and DL are supported by grants from the NIH: DP5-OD021396, R01 AI136680, and U54 GM103368.

### Appendix A. Some details for calculations in Section 3: geometrical factor for decay of density Eq (3.5), checking numerical sampling code Eq (3.9), creating distributions according to some prescribed function Eq (3.15), proof that uniform distribution maximizes the SCF Eq (3.22), derivation of Eq. (3.23) for the SCF for modulated side-views

#### A.1 A general formula for the projection geometrical factor: Eq. (3.5)

Our claim in Eq (3.5) is that

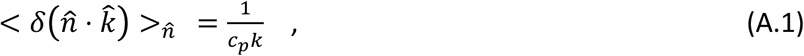

where *C*_*p*_ is a geometrical factor that we wish to calculate in general dimensions, especially for *D* = 2, 3. By 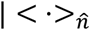, we mean the average over the surface of the until ball in D dimensions. One easy way is to integrate the above equation over all 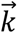 with *k* < *L* in D dimensions. Then on the left-hand side we get:

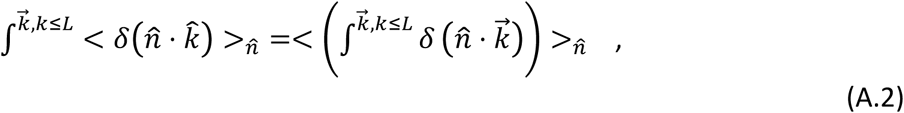

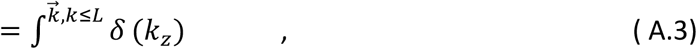

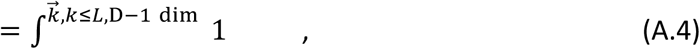

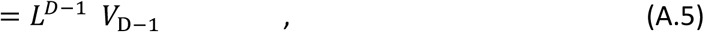

where *V*_D−1_ is the volume of the unit ball in *D* − 1 dimensions. Eq (A.3) holds because the integrand in (A.2) is no longer dependent on the direction, 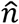, so the average over 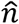 seen in (A.2) integrates to 1. Moreover 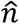 where it appears in the integral may be set to 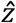 for convenience. On the RHS of (A.1) we also perform the integration over the ball of radius *L* and get

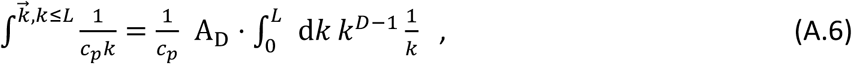

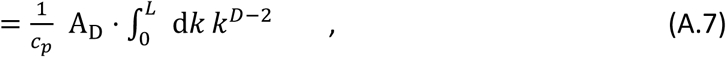

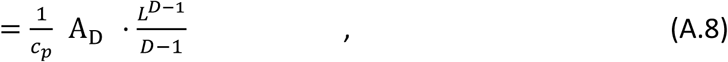

where *A*_D_ is the surface area of the unit ball in *D* dimensions.

Equating the last two expressions shows that

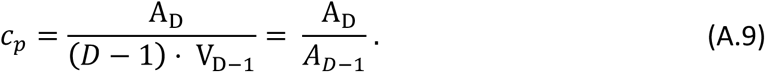

This gives

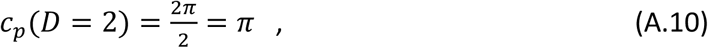

and

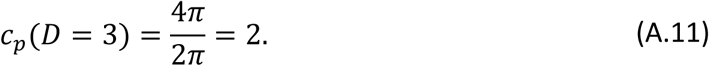

Therefore, the geometrical factor 2 that appears in Eq 3.5 is simply the ratio of the surface area of a unit ball to the circumference of a great circle of the same ball.

#### A.2 Checking the Sampling Code Eq. (3.9)

We want to evaluate

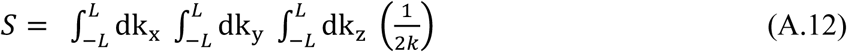

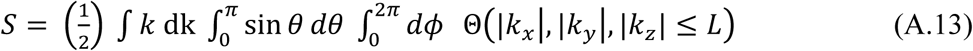

It is enough to consider the upper quadrant, where all the components are positive.

This is where both the azimuthal angle, *ϕ*, and the spherical angle, *θ*, are in the range 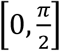. This gives us a symmetrization factor of 8. However, we may also consider a definite ordering for the *k*_*x*_, *k*_*y*_, *k*_*z*_ giving a symmetrization factor of 6. Putting this together we have

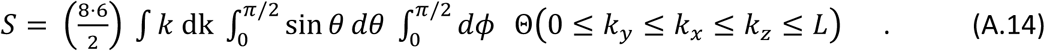

We wish to reorder the integrations: first *k*, then *θ* then *ϕ*. The spherical representations for the components may be written as: *k*_*x*_, *k*_*y*_, *k*_*z*_ = sin *θ* cos *ϕ*, sin *θ* sin *ϕ*, cos *θ*)

Now 0 ≤ *k*_*y*_ ≤ *k*_*x*_ is easily represented by 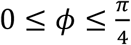. Let’s write down what we have so far:

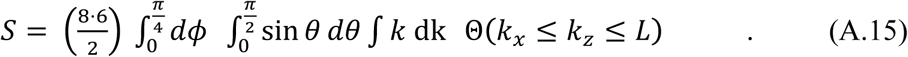

To ensure the last two inequalities we need *k* ≤ *L*/ cos *θ* and tan *θ* cos *ϕ* ≤ 1. This last inequality can be used to govern the upper limit of the *θ* integration, in place of the *π*/2, because tan *θ* can always attain the value 1/ cos *ϕ* on the interval [0, *π*/2].

Putting this all together and developing we get

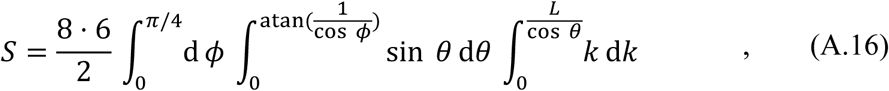

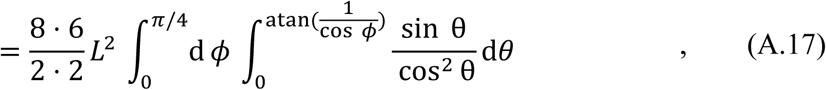

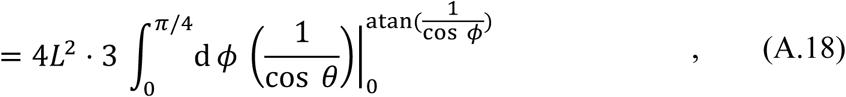

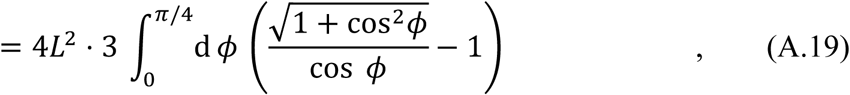

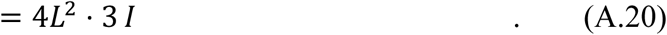

To evaluate the last integral, *I* we use the substitution 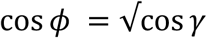. The limits for *γ* now become *ϕ* = 0 ↔ *γ* = 0, *ϕ* = *π*/4 ↔ *γ* = *π*/3. We also have 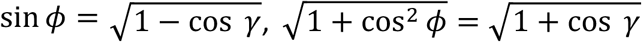. This leads to 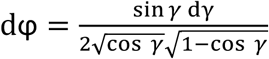, which can be simplified to 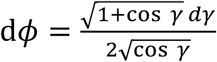. So

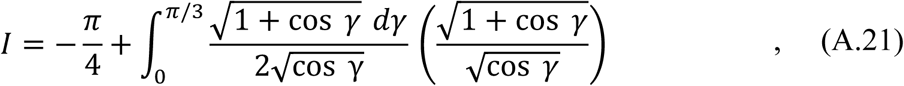

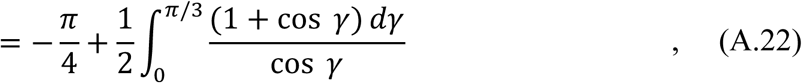

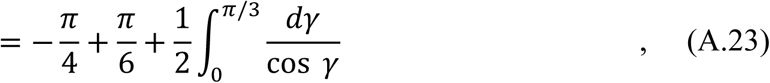

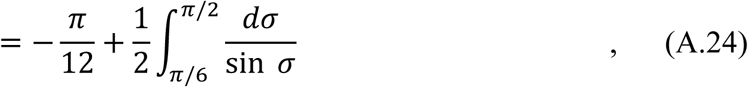

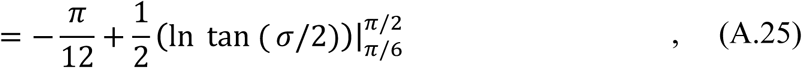

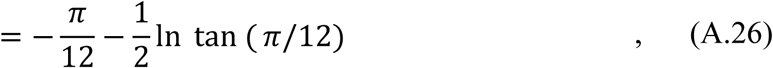

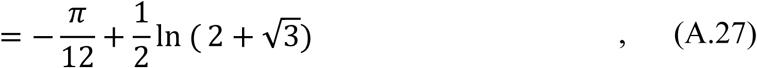

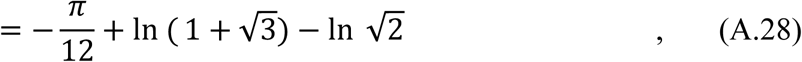

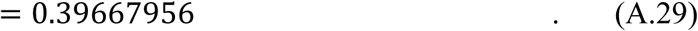

Finally, now

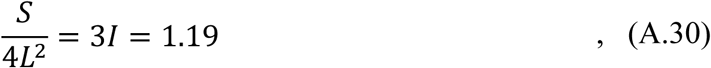

This factor is the empirically observed excess area of an average plane embedded into a cube. This is the approximately 20 per cent increase in actively sampled points. It is a larger factor than the comparable 1.12 that would appear in a similar 2D problem.

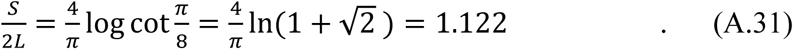

Another approach to evaluating (A.9) is to introduce an auxiliary variable via 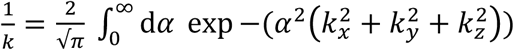. Then there are just a few steps to a single integral and a numerical evaluation: 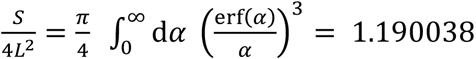, which is (A.30).

#### A.3 Creating distributions according to some prescribed function Eq (3.15)

In order to create a numerical sampling map for modulated side views, we would like to assign azimuthal angles to projections such that the oscillatory azimuthal distribution density indicated by (3.11) is achieved. This is well known how to do: for completeness, we include the argument here. From the density function (3.11), the cumulative distribution function can be found which is

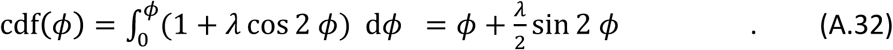

Now the azimuthal angle should be given by

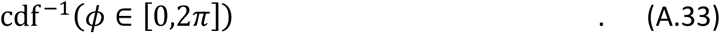

That is, numbers should be drawn evenly between 0 and 2*π*, resulting in an array given by (A.33). These are the angle labels to be given to achieve the desired distribution (3.11). So long as *λ* < 1, this is easy to do, because the distribution is positive and the cdf is monotonically increasing (graphically, the inverse corresponds to flipping across the diagonal, which maps a function into another function due to monotonicity). The python pseudo code would read: phi0= cdf0= np.linspace(0,2*np.pi,NumPoints); 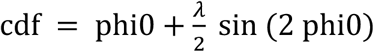, cdfInv = np.interp(phi0,cdf, cdf0). That is, map the array phi0 to the desired phi (which is the desired cdfInv), in the same manner that cdf was mapped to cdf0, where phi0, cdf0 are both regularly spaced.

#### A.4 Proof that uniform distribution maximizes the SCF Eq (3.22)

Consider a set of positive numbers {*α*_*i*_} that satisfy a constraint *C*: **Σ**_*i*_ *a*_*i*_ = *M*. The set are to represent the sampling on the unit sphere. We wish to maximize 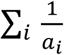. We begin by writing the usual variational:

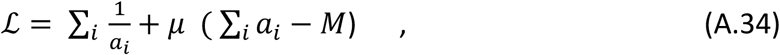

where *µ* is a Lagrange parameter. Extremizing *ℒ* wrt the *a*_*j*_ yields

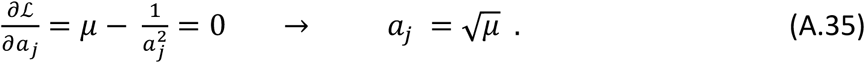

The second variation is:

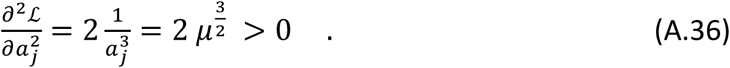

Since the second variation is positive, the uniform solution *a*_*j*_ = constant, corresponds to a minimum.

The argument supplied here implies why the SCF attains its maximum (1/SCF attains its minimum as in the above calculation), when the sampling (which is a conserved quantity on every shell of Fourier space) is distributed as uniformly as possible, or equivalently the projections are distributed uniformly.

#### A.5 Derivation of Eq. (3.23); SCF for modulated side-views

From Eq. 3.16, we have

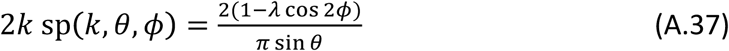

Using the definition of SCF, 1/SCF = < (1/(2k sp)>, then (A.37) becomes

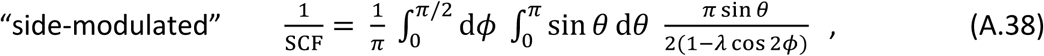

where the last term in the integrand of (A.38) is the reciprocal of (A.37). The integration over *θ*, can be easily performed 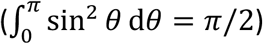 leaving:

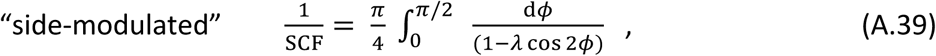

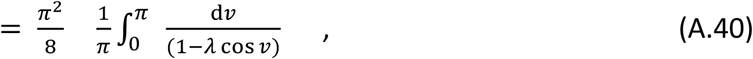

Integrals of the sort that appear in (A.40) are easily reduced by means of the so-called Weierstrass half angle formula: 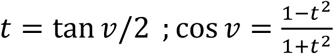. The integral in (A.40) becomes 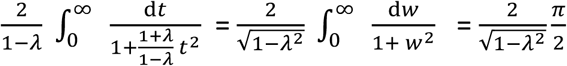. So the expression in (A.40) becomes:

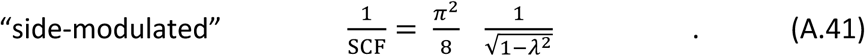

which is (3.23).

### Appendix B Derivation of Eq. (3.12) and (3.18): sampling distributions from projection distributions

Eq (3.11) reads

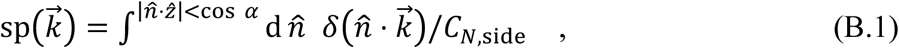

where integrations are taken over all unit vectors, 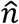, in 3D. Also *C*_*N*,side_ is a normalization constant ensuring Eq (3.8): 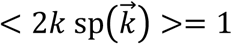, where 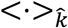 denotes angular average over the angles in *k* with the uniform measure on the sphere. The integration in B.1 is over the set of normal vectors to the sphere, with the given constraint. Putting this together with B.1 yields:

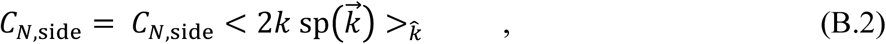

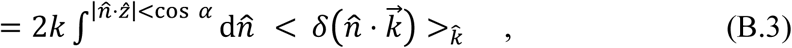

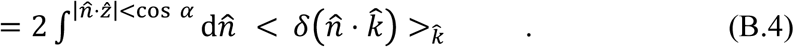

Now 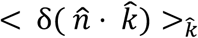 cannot be a function of the direction of 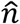. So it can be conveniently calculated when 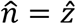, which does not depend on an azimuthal angle in the integration over 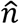, and therefore leads only to the average over the altitude. This leads to:

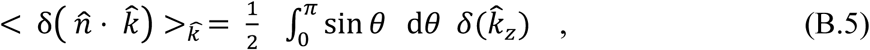

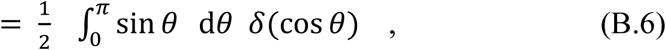

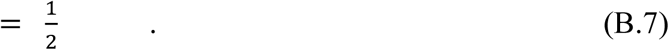

Eq. B.7 is a natural result. It is the ratio of the circumference to the surface area of the unit circle: 2*π*/4*π* = 1/2. Returning to (B.4) we get:

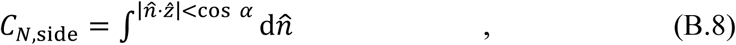

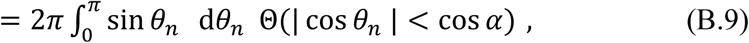

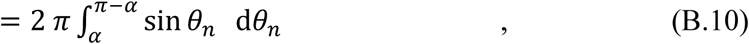

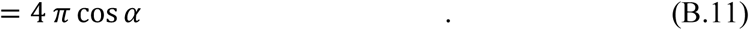

So, substituting (B.11) into (B.1) yields

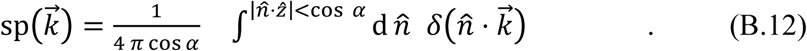

It is easy to argue that 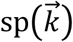 does not depend on the azimuthal angle of 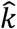, which we can therefore take to be zero in order to evaluate (B.12): 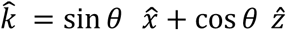. Instead of the integration over the sphere given by the unit vector, 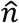, we need to perform the integral in (B.12) over the great circle perpendicular to 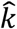. Therefore, we can parametrize 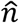, in the integration in (B.12) by

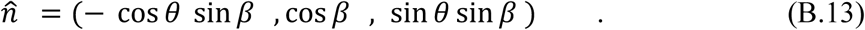

Eq. (B.13) is a parametrization of all the unit vectors perpendicular to 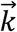 as described in the last paragraph. By changing *β*, we can sweep out the unit vector given by (B.13): these are the locus of normals to 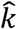 and outside the cone of half angle *α*. So from (B.12)

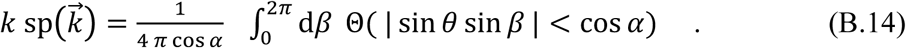

The criterion Θ (| sin *θ* sin *β* | < cos *α*) in B.14 is a rewrite for the constraint of the projection directions, 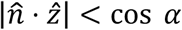, from Eq. B.12. Continuing from Eq. B.14.

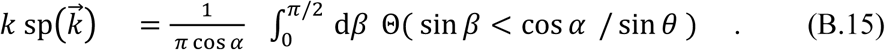

If cos *α* > sin *θ*, then the argument of the indicator function in (B.15) is always true. If not the upper limit of *β* in the integral must be reduced to asin (cos *α* / sin *θ*). This leads to:

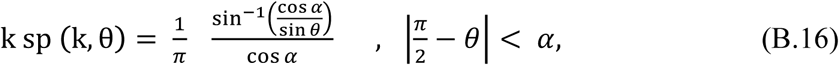

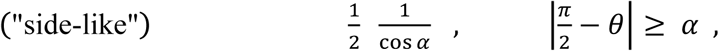

which is (3.12).

Finally

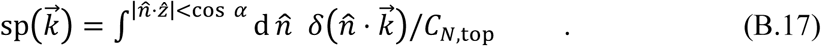

Using (B.8) and (B.17) using the parallel argument to (B.1)-(B.7) together, we note that

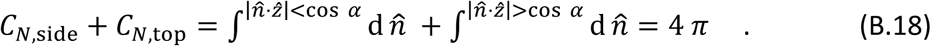

So

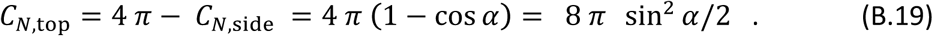

The parallel derivation to (B.14) now becomes:

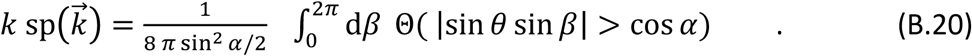

This is the integration around the locus of points normal to 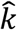 and inside the cone of half-angle, *α*. However, sin *β* may be replaced by cos *β* by shift of origin, and an overall factor of 4 introduced due to the 4 equivalent quadrants:

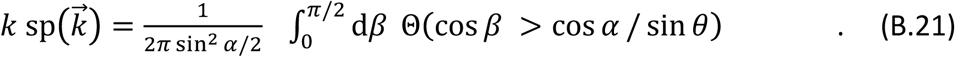

If cos *α* > sin *θ*, then the condition of the indicator function cannot be fulfilled, and the left-hand side = 0. Otherwise

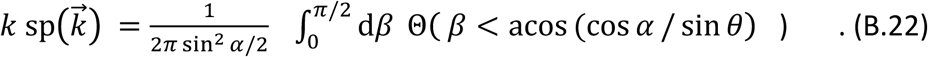

So

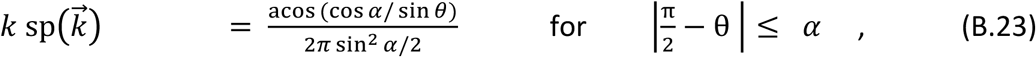

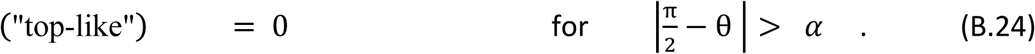

This is (3.18). Thus, the sampling is zero in directions close to along the z-axis, for the top like cases.

## Glossary

FSC(*k*): is the Fourier shell correlation of the reconstruction at Fourier frequency *k*.
SSNR (*k*): is the spectral signal to noise ratio of the reconstruction at Fourier frequency *k*.
ssnr: (*k*) is the per particle SSNR, used in the discussion in Section 4.
*L*: the side of the real space box
*N*: the number of particles in the reconstruction
*k*: is Fourier magnitude
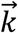: is a 2D or 3D point in Fourier space
SCF: is the sampling compensation factor, characterizes effect of sampling on SSNR
𝒩 (*k*): is noise-to-signal power (sections 1,2)
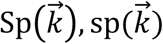: is the sampling function (and per particle sampling function) defined at a 3D lattice site 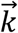
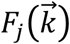: is the Fourier value of the j^th^ projection at the point 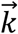.
*M*_*j*_(*k*): is the effect of the microscope (CTF) on the *j*^th^ projection
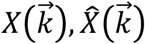: is the target model and a running estimate of the model
*R*_*j*_: is a 3D rotation matrix describing the projection, *j*.
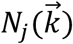: is the noise added to the projection, *j*.
*E*(*k*): is the total envelope that attenuate the image due to microscope and misalignment effects.
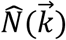: is the effective noise at 3D lattice sites after regrouping from projections
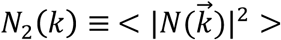: is the power of the noise
*F, G*: are half maps used to derive FSC relations in section 2
*P*_*k*_, *Q*_*k*_: the number of measured and unmeasured voxels, on a Fourier shell of radius *k*, when there is missing data.
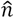: is used as a unit vector demarcating a projection
Θ (x): indicator function, which is 1 if the condition x is true, 0 otherwise
*λ*: is the amplitude of the modulation for the modulation of side views (section 3)
*α*: is a cone half angle: for top-like views, projections are inside cone; for side like, projections are outside the cone.
*ϵ*: is the fraction of projections that are not restricted to be in the main cone of half angle *α* (Section 3)

Euler Angle is one of the three angles used to describe rotation matrices (*θ* is rotation around Z-axis, *ϕ* is rotation around Y-axis, *ψ* is in-plane rotation).

